# Large-scale exploration of protein space by automated NMR

**DOI:** 10.64898/2026.02.16.706194

**Authors:** Thomas Müntener, Dylan Abramson, Elsa Stern, Ines Hertel, Gytis Jankevicius, Guillaume Mas, Gert E. Folkers, Basile I. M. Wicky, Sebastian Hiller

**Author notes:** equal contribution (Thomas Müntener, Dylan Abramson).

## Abstract

Protein structures can now be predicted and designed at scale, yet experimental access to dynamics and conformational heterogeneity remains limited in throughput. This gap prevents a systematic understanding of how protein sequences encode motion and functional flexibility. Here, we establish a scalable experimental pipeline combining protein design, automated production, and nuclear magnetic resonance (NMR) spectroscopy to enable high-throughput characterization of protein structure and dynamics at atomic resolution. A single operator can produce and analyze hundreds of isotopically labeled proteins per week, with per-sample cost largely defined by DNA synthesis. To benchmark this approach, we experimentally characterized 384 de novo designed proteins spanning diverse regions of structure space. High-quality two-dimensional NMR spectra were obtained for 239 samples (62% of designs overall). NMR characterization confirmed that the designed proteins adopt their intended folds, and revealed unexpected local dynamics that are not captured by current computational models. Our approach establishes a foundation for data-driven modelling of sequence–structure–dynamics relationships and unlocks a new regime of statistical structural biology, where insight into protein biophysics is gained from experimental ensemble studies of suitably designed protein clusters.

## Introduction

Understanding how amino-acid sequences encode protein structure, dynamics, and function is a central goal of structural biology^1^. Fueled by the availability of large datasets of protein structures, recent advances in deep learning have transformed our ability to predict and design proteins with atomic accuracy, enabling exploration of vast regions of protein space *in silico*^2–6^. These advances have primarily addressed the mapping of sequence to static structures. Experimental characterization of protein dynamics, disordered regions, conformational heterogeneity, and folding behavior remains fundamentally limited in scale, preventing systematic investigation of how protein sequences encode energy landscapes,motion, and functional flexibility.

Nuclear magnetic resonance (NMR) spectroscopy provides uniquely rich experimental access to protein structure and dynamics across multiple timescales at atomic resolution.^7–9^ Yet, despite decades of methodological development, NMR has largely remained a low-throughput technique, typically applied to individual proteins or small sets of related targets^10^. Large-scale structural genomics initiatives have demonstrated that systematic structure determination is possible^11–13^, but at substantial financial and logistical cost, and they did not address the broader challenge of generating standardized datasets suitable for the application of deep learning techniques. Notable approaches to parallelize expression were made but have not found widespread use^14–17^. So, while structural databases have enabled machine learning approaches for protein structure prediction, the NMR field lacks standardized datasets at the scale that would enable deep learning approaches for assignment, function, dynamics, and conformational heterogeneity. De novo protein design offers an attractive route to generate such datasets: generative models can systematically sample structure space, and designed proteins typically express robustly^18–20^, making them ideally-suited for high-throughput production. Combined with automated NMR characterization, this approach could enable statistical analysis of protein behavior across large cohorts, analogous to how structural databases have enabled data-driven advances in structure prediction.

Here, we establish a scalable experimental platform that integrates generative protein design, automated high-throughput protein production, and standardized NMR spectroscopy to enable atomic-resolution characterization of protein structure and dynamics across hundreds of proteins. We demonstrate feasibility on a library of 384 de novo designed proteins spanning diverse regions of structure space, obtaining high-quality spectra for 62% of samples. Our platform thus establishes a foundation for data-driven modelling of sequence–structure–dynamics relationships and unlocks a new regime of statistical structural biology, where insights on biophysical properties, mechanisms, and functions are gained from experimental studies on ensembles of suitably designed proteins.

## Results

### Design of a structurally diverse protein library

To ensure diversity and broad representation of structure space we leveraged protein design to generate samples de novo, employing two different models for the task: RFdiffusion^5^, a common choice for protein backbone generation, and Proteína^21^, a newer-generation model based on flow-matching. RFdiffusion has been extensively validated experimentally, but tends to produce simple helical structures^22^, whereas Proteína, though lacking prior experimental validation, has been shown to produce more diverse structures.

Designs were created using a canonical structure-based design pipeline, starting with backbone generation, followed by sequence design with ProteinMPNN^18^ and refolding with Boltz-1^23^ (**Figure 1**). For each model, 17,500 backbones were generated, and 10 sequences per backbone were designed. All designed sequences were refolded *in silico*, and filtered based on structural self-consistency and confidence metrics (see Methods for details). Because simple helical structures tend to score better, we clustered the designs using Foldseek^24^, and sampled from each cluster to ensure broad coverage of structure space (**Figure 2A**). As expected for designed proteins, local sequence-structure relationships were idealized, resulting in a high content of glutamate residues, followed by lysines and leucines, which together accounted for, on average, 40% of the designed sequences (**Figure S1**). A higher content of β-sheet structures correlated with an increased number of glycines, valines and threonines in the sequence.

**Figure 1.**
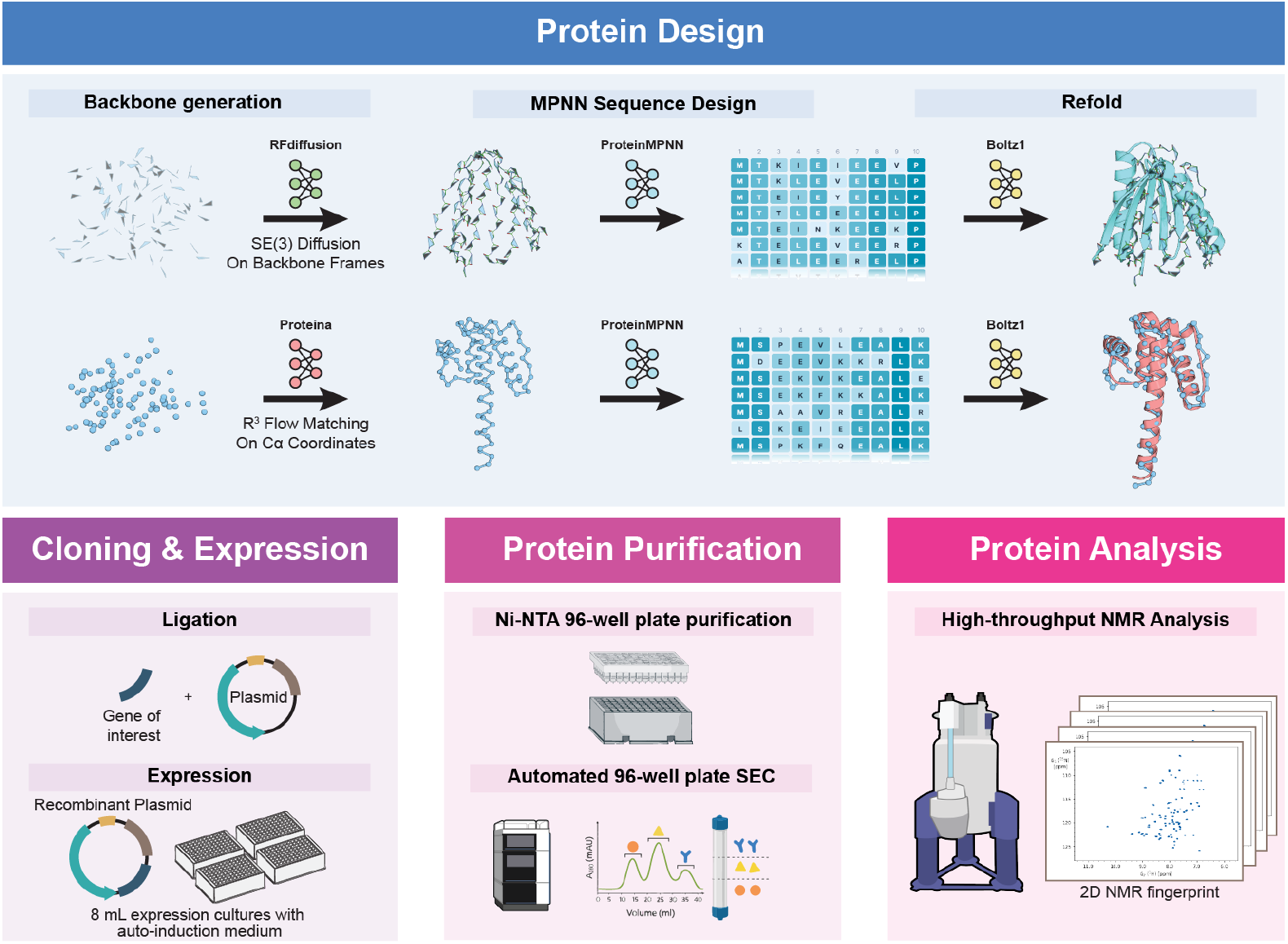
Protein design and automated production workflow. Design pipeline (top); de novo backbones were generated using either RFdiffusion, or Proteína, followed by sequence design with ProteinMPNN and filtering with Boltz-1. Experimental validation (bottom); designs were ordered as synthetic genes, cloned into expression vectors, and produced in *E. coli* from ^15^N-labelled autoinduction media in 96-deepwell plates (left). Samples were purified by immobilized metal affinity chromatography in 96-well plate format, followed by high-throughput size-exclusion chromatography (middle). NMR fingerprints are obtained in 45 min for all produced samples.

**Figure 2.**
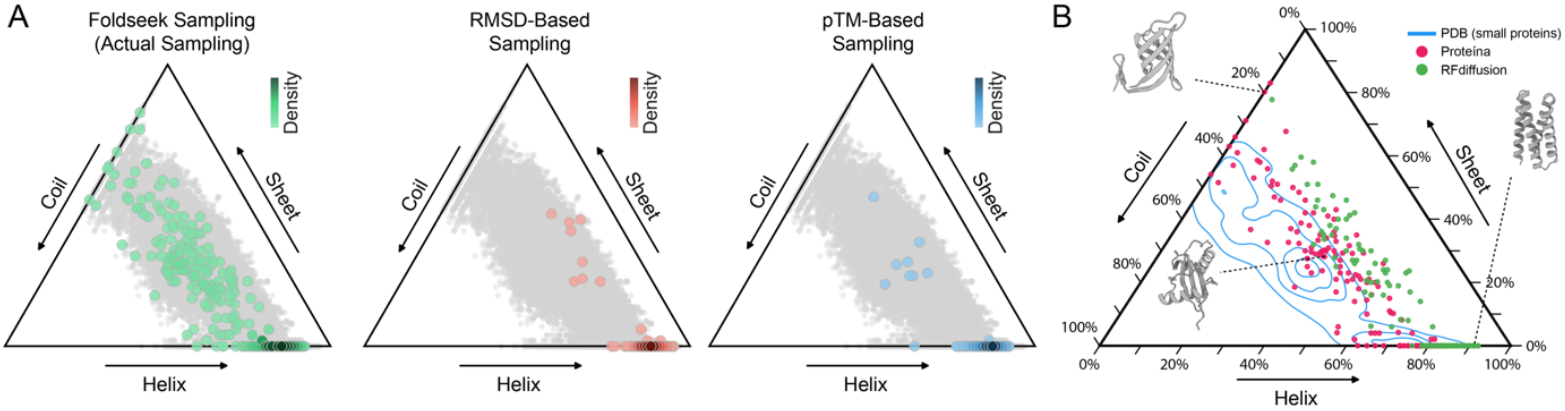
Large-scale coverage of structural space using de novo protein design. (**A**) Impact of sampling strategy on structural coverage. Grey dots show all *in silico* successful designs while colored dots show samples that would be picked for experimental testing. Foldseek clustering ensures adequate representation of the structural space (left), while sampling purely on quality metrics (RMSD-based, pTM-based, middle, right) generates distributions heavily skewed towards helical structures. (**B**) Ternary plot showing the distribution of secondary structure content for designed proteins (Proteína: n = 192; RFdiffusion: n = 192), compared to structures of similar size deposited in the PDB (n = 3650, contour lines representing kernel density estimates).

In total, 384 designs were selected for experimental characterization, 192 for each of the two backbone generation methods. While α-helices remained the dominant structural element present in the designs, a substantial amount of folds (more than 30%) contained β-sheet elements, and some were even completely devoid of α-helices. This library thus constitutes a synthetic mimetic of natural proteins of similar size deposited in the PDB (**Figure 2B**).

### High-throughput protein production

For expression of the proteins in isotopically labelled form, we adapted the protocol from Qian *et al*^25^. Designs were codon-optimized for expression in *E. coli* and ordered as synthetic DNA. Gene fragments were cloned into expression vectors by Golden Gate Assembly using an acoustic liquid robot. Final constructs contained a C-terminal SNAC-tag^26^ followed by a His_6_-tag, both of which were kept during the subsequent NMR analysis. Constructs were directly transformed into *E. coli* expression strains, and proteins were expressed in 96-deepwell plates by auto-induction using ^15^N-isotopical labelling. Purification followed a two-step process, starting with chemical lysis and IMAC purification in 96-well plates, followed by automated semi-preparative size-exclusion chromatography. Automated analysis scripts performed sample quality control and quantification from the resulting chromatograms, as well as estimation of the protein oligomeric state from the elution profiles. Fractions were selected automatically from the elution profiles and pooled by a liquid handling robot, resulting in a median yield of 115 μg per sample. Of the 384 selected designs, only 5 samples failed during initial expression and purification (1.3%), while the remaining 379 proteins (98.7%) were advanced to NMR analysis. The combination of the favorable manufacturability of designed proteins with automated protein expression and purification effectively removes sample production as a bottleneck for high-throughput NMR spectroscopy.

### Automated NMR screening

We applied a standardized NMR screening pipeline to all 379 proteins, recording 2D [^15^N,_1_H]-HMQC experiments at 25 °C for 45 minutes. The resulting 2D NMR spectrum is commonly used as a protein fingerprint, providing a direct assessment of protein quality and, through line shapes, initial insight into protein dynamics. Importantly, once samples were loaded, data acquisition proceeded without manual intervention, making spectrometer time — rather than personnel — the limiting factor of this step. At this rate, a single spectrometer characterizes 32 samples per day, or 224 per week. Analysis of spectra quality was automated, based on the number and intensity distribution of peaks (see Methods for details). Of the 379 proteins, 239 samples (62% of designs) generated high-quality spectra suitable for detailed analysis (**Figure 3, Figure S2**). 79 samples (21% of designs) generated spectra with detectable peaks, these were however characterized by many missing peaks and a low signal-to-noise ratio and thus of low quality. The remaining 61 samples (16% of designs) did not yield interpretable NMR spectra within the 45 min acquisition time. Plausible explanations are low expression, the presence of aggregates, and multimerization. The quality of the spectra strongly correlated with the protein concentration in the measured NMR sample as expected (**Figure 3C**). Most good/very good spectra were obtained from samples with protein concentration above 10 µM. Among these, the success rate was 87%. We note that by simply doubling expression volumes to 16 mL, which can readily be done in future runs, the resulting 4-fold speedup in acquisition time would enable a throughput close to a thousand spectra per week.

**Figure 3.**
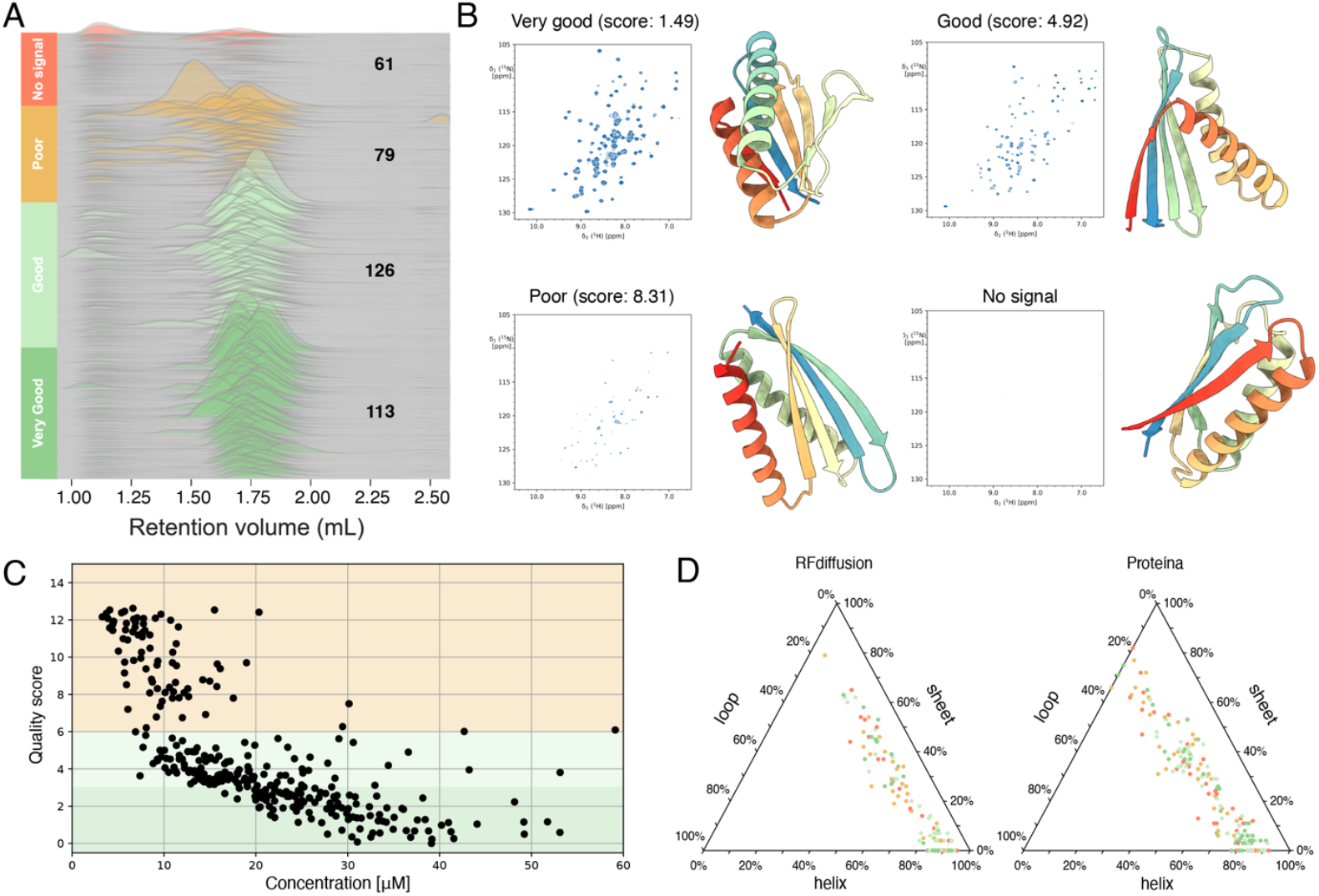
High-throughput NMR analysis of de novo designed proteins. (**A**) SEC profiles for the 379 expressed designs. Monomeric proteins of these sizes are expected to elute ∼1.75 mL on this column (Superdex 75 5/150). Counts are given for each NMR quality class. (**B**) Examples of four designer proteins and their 2D [^15^N,^1^H]-SOFAST HMQC NMR spectrum. Proteins were produced in 96-well plate format, and NMR spectra were recorded at 25°C in 45 minutes on a 600 MHz spectrometer. (**C**) NMR quality score as a function of sample concentration for 318 designer proteins. (**D**) Ternary plots of predicted secondary-structure composition for proteins designed with RFdiffusion (left) and Proteína (right). Each point represents one designed protein positioned by its helix, sheet, and loop fractions. Colors indicate the experimental spectrum quality as defined in A). In both cases, a large fraction of the density is located at the bottom right corner of the plot (high helical content, cf. Figure. 2).

### Statistical analysis of 2D NMR fingerprints

We performed a detailed analysis of the large 2D NMR spectra dataset to identify correlations between sequence features, spectral quality, and structural characteristics. The two computational models Proteína and RFdiffusion yielded similar numbers of proteins with NMR spectral quality classified as “Good” or “Very good”. Purely helical proteins featured similar experimental success rates (70%) with either computational model, whereas with increasing β-sheet content, the more recent model, Proteína, showed a greater experimental success rate (**Figure S3**).

To explore whether computational metrics could predict experimental outcomes, we calculated aggregate observables from HMQC spectra and correlated these with *in silico* features derived from the designed models. Since the majority of HMQC spectra were unassigned, we looked at whole-spectrum metrics. The strongest correlations (R^2^ up to 0.76, **Figure S4**) were observed for mean chemical shifts, which recapitulate secondary structure composition. The data show that the average of both the ^1^H and _15_N chemical shifts increases with the percentage of β-sheet in the protein (**Figure S5**), in full agreement with established knowledge^27,28^. This finding provides an internal consistency check that at least at the ensemble level, the designed proteins adopt their intended conformations. Unsupervised clustering of spectra showed clear separation between fold classes (**Figure 4**), suggesting that the application of machine learning techniques on large-scale systematic NMR data can uncover patterns in unassigned spectra.

**Figure 4.**
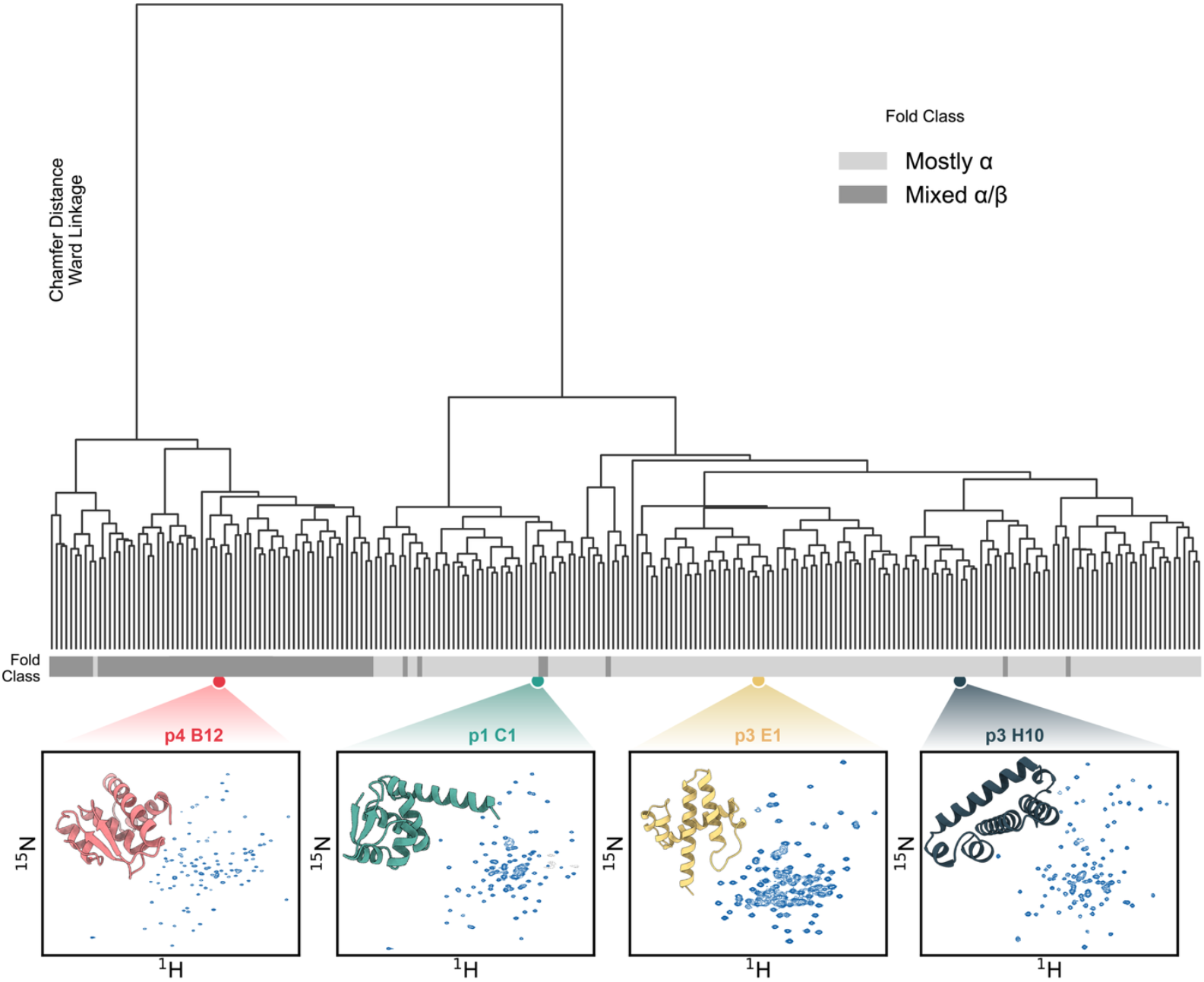
Similarity analysis of 2D NMR spectra. Dendrogram, where each leaf represents one 2D [^15^N,^1^H]-HMQC spectrum of the 239 designer proteins with good /very good quality. The annotation bar beneath the dendrogram shows secondary-structure content. HMQC spectra from designs with predominantly helical folds form a cluster clearly distinct from those with mixed α/β content. Spectra and structures are shown for four representative designs.

Beyond this, we sought metrics that could report on conformational dynamics. The coefficient of variation (CV) of peak intensities emerged as the most informative dynamics-related observable, showing moderate correlations with computational features with an R^2^ up to 0.23 for some parameters (**Figure S4**). CV captures the heterogeneity of the spectral fingerprint with higher values indicating greater intensity dispersion. This variability can reflect conformational exchange, differential relaxation rates, or local unfolding. The CV is concentration independent and thus self-normalizing and this may explain its robustness compared to other intensity-based metrics. Together, the detection of these correlations proves that structural and dynamic protein features can be recovered from statistical analyses of experimental ensemble data.

### Structure validation

From the full dataset, we selected 9 proteins for advanced characterization. Each of these proteins was re-expressed with ^13^C-labeling in a larger expression volume. The proteins were then subjected to sequence-specific resonance assignment using an HNCACB experiment and automated FLYA/CYANA^29^ assignment from manually curated peak lists, requiring about one day of experiment time and three hours of manual work per protein. For six proteins, almost complete assignment was achieved directly, and ambiguities for the three remaining proteins were resolved by using an additional 4D APSY-HNCOCA (oaNH). With the assignment at hand, secondary chemical shift analysis was used to determine the type and location of secondary structure elements (**Figure S6**). For all nine proteins, these were matching precisely with the expectations from the designed model, indicative of correct folding.

We then aimed to validate one representative protein to full extent and determined its NMR structure (**Figure 5**), using a comprehensive set of backbone and side-chain experiments, complemented by NOESY spectra for distance restraints. The resulting structure was in excellent agreement with the predicted model, with RMSD to the designed model of 1.3 Å. This protein exhibited a second minor conformation, observable by a set of peaks indicative of a second state, most clearly visible for residues 88–92 (**Figure 5E**). This second state is most likely due to a cis/trans isomerization of the proline residue at position 82. Second states were also observed in several other proteins upon close inspection (**Figure S7**). Some of these minor states correspond to only 5% of the intensity of the major state. The observation of lowly populated states in several designs demonstrates that our platform has sufficient sensitivity and dynamic range to systematically study designed multi-state proteins.

**Figure 5.**
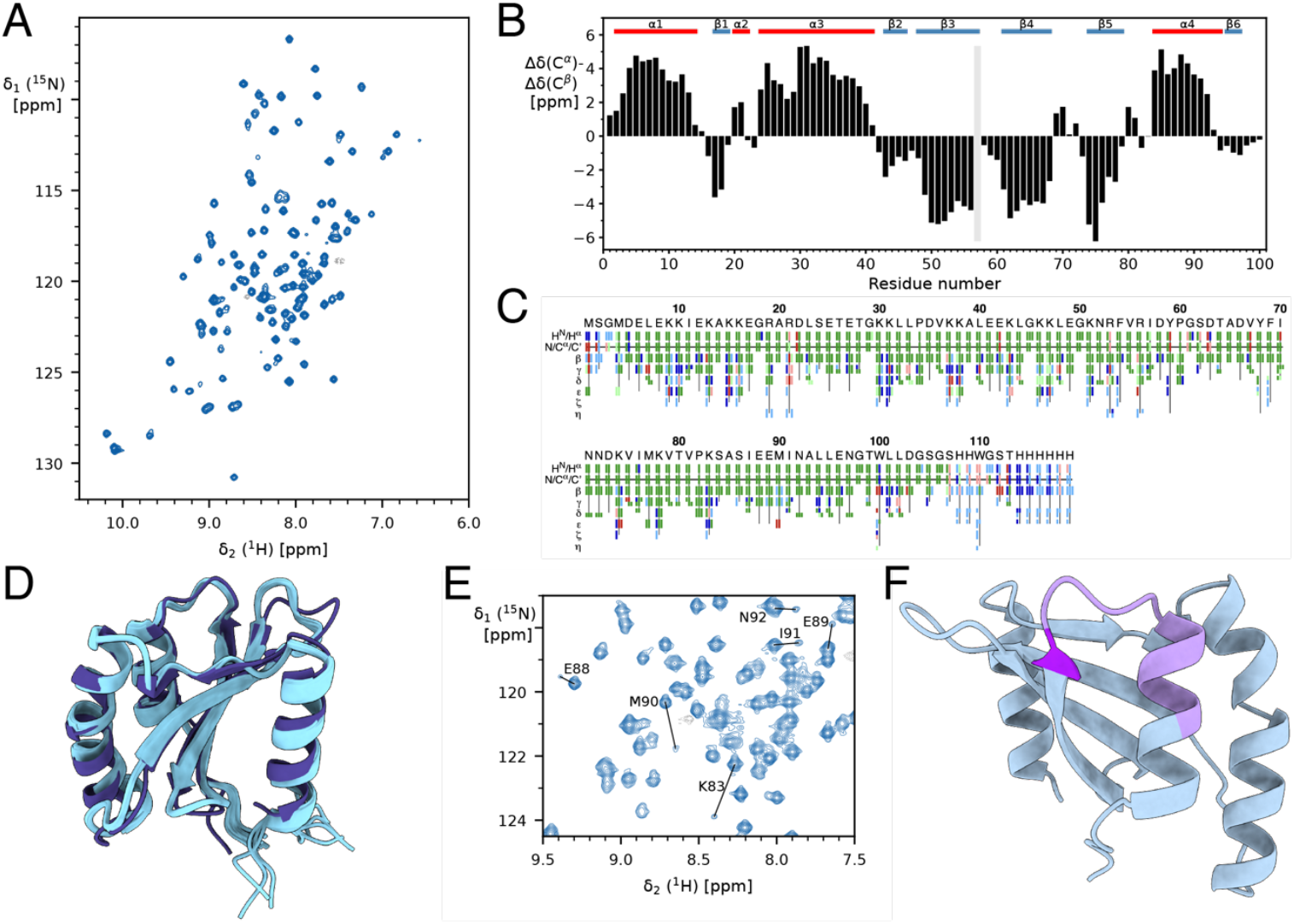
Structural characterization of designed protein p2 A4. (**A**) 2D [^15^N,^1^H]-SOFAST-HMQC spectrum showing good chemical-shift dispersion consistent with a folded state. (**B**) Secondary C_α_/C_β_ chemical-shift deviations relative to random coil, reporting on secondary structure (positive values indicate α-helical propensity; negative values indicate β-sheet propensity). The designed secondary-structure pattern is indicated above. (**C**) Summary output of the automated resonance assignment generated with CYANA. (**D**) Superposition of the NMR-derived structural ensemble (10 structures) for p2 A4 (light blue) with the designed structure (dark blue). (**E**) Identification of a second conformation caused by cis/trans proline isomerization at residue 82 spanning residues 83 and 88–92. (**F**) Structure of protein p2 A4 highlighting proline residue 82 in dark purple and residues affected by its isomerization in light purple.

### Protein dynamics

We recorded NMR relaxation experiments to characterize backbone dynamics and conformational flexibility of the 9 proteins with available backbone assignment (**Figure 6**). Importantly, these experiments were recorded on the samples resulting from the high-throughput production scheme, i.e. with protein concentrations in the range 10–50 μM. For each sample, the experiment time was 32 h. Several of the proteins showed non-uniform local dynamics, such as loop or segments of secondary structure motions, which is intriguing as the proteins had been designed as static entities. Most interestingly, some loops showed clear presence of local dynamics, while other similar loops did not. It has been reported that de novo designs exhibit frustrated energy landscapes, occupying intermediate local minima rather than cooperative folding transitions^30–33^. This frustration can manifest as persistent backbone dynamics in regions that would typically be rigid in natural proteins, and it will be interesting to explore these connections better in future experiments.

**Figure 6.**
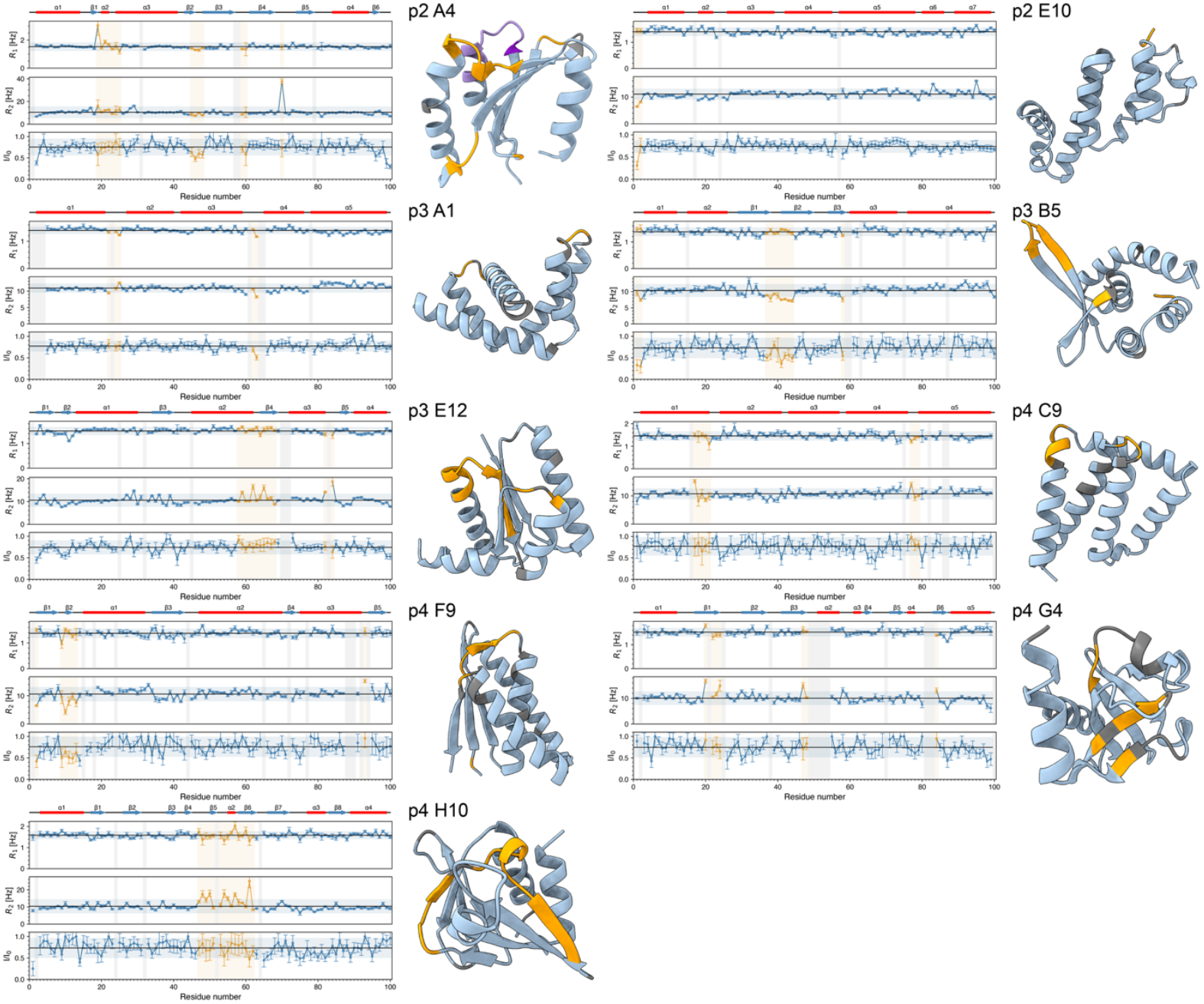
Backbone dynamics of nine selected designer proteins. Residue-specific *R*_1_(^15^N), *R*_2_(^15^N), and {^1^H}–^15^N NOE data were recorded at 25°C on a 600 MHz spectrometer. Unassigned residues are shown in grey. Segments with increased local dynamics are colored yellow, as evidenced by *R*_2_, *R*_1_ and/or {^1^H}–^15^N NOE significantly deviating from the global average. Residues affected by a *cis*/*trans* isomerization are colored in purple.

When we examined residue-level computational predictions with NMR relaxation parameters across the 9 designs with backbone assignments, we found essentially no correlation (R^2^ < 0.12 for all comparisons, **Figure S8**), suggesting that learned structural priors from structure prediction networks may be insufficient to capture residue-specific dynamics in designed proteins. These observations highlight the gap between generative models trained on static structures and experimental behavior, indicating the need for our platform to systematically explore the dynamics of designer proteins.

## Discussion

Here we show that modern protein design, combined with automated protein production and NMR analysis, can deliver atomic-resolution data on structurally diverse proteins in a high-throughput, low-cost manner (about 25 USD/sample, dominated by the cost of synthetic DNA). We term this approach NMR-Automated Protein Production (NMR-APP), which enables a single operator to produce data at the rate of NMR data acquisition, corresponding to, e.g., 32 proteins per day for 2D fingerprint spectra.

While simple structures might now be trivially obtained from sequence using deep learning architectures such as AlphaFold, our results reveal that computational metrics provide a weak indication about the dynamics of designed proteins. While recent work showed that pLDDT moderately correlates with order parameters for natural proteins^34,35^, successful dynamics prediction for natural proteins may rely critically on evolutionary information. The fact that recent protein language models can predict dynamics from sequence alone supports this idea^36^. This suggests that the ability of these models to predict protein motion for natural sequences may not be based on learned priors of protein dynamics, but rather on statistical patterns extracted from evolutionary data. This limitation motivates precisely the type of large-scale, standardized NMR datasets that our pipeline enables: datasets that could eventually train models to predict dynamics from first principles, independent of evolutionary information.

In future implementations, the NMR-APP platform can be expanded to include side-chain-specific experiments, such as specific methyl labeling on deuterated background^37^, enabling its application to increasingly larger proteins and more complex systems. While we leveraged the favorable manufacturing properties of designed proteins for this work, we note that the experimental pipeline itself is not restricted to de novo sequences, and can be extended to natural proteins and complexes using appropriate labeling strategies.

Sequence-specific resonance assignment remains a key bottleneck of NMR spectroscopy workflows, including NMR-APP. In the current implementation, assignment of individual proteins required additional spectroscopy and semi-manual assignment procedures. These procedures will need to be further time-optimized by determining the minimal set of spectra needed for automated assignment, such as down-sampled NOESY or other rapidly sampled, high-complexity spectra^38^. With the accumulation of sufficiently large datasets, we expect that advanced automated assignment will become feasible through advanced machine learning methods and that the minimally required experimental input will decrease as these methods advance. Because the protein structure is known, it might well be possible in the future to infer the sequence-specific resonance assignment from the 2D HMQC fingerprint spectrum directly.

Our results break a long-standing throughput barrier in NMR structural biology, improving the time and costs required for data collection by orders of magnitude compared to previous structural proteomics efforts. The NMR-APP approach enables the transition from single-protein structural biology to statistical structural biology, an experimental regime where insights are gained from large-scale experiments on ensembles of suitably designed proteins. Data that previously required years of effort can now be generated in a matter of weeks. Such an NMR-driven statistical structural biology will provide new insights into protein dynamics, multi-state behavior, allostery, functional heterogeneity, intrinsically disordered proteins, energy landscapes, and more.

## Acknowledgements

SH and BIMW acknowledge generous core funding from the University of Basel and the ETH Zurich, respectively. DA is supported by funding from the National Center of Competence in Research (NCCR) Molecular Systems Engineering (MSE). Large-scale design and structure predictions were performed on the Euler cluster of ETH Zurich.

## Materials and Methods

### Protein design

For both models, 17,500 initial backbones were generated and 10 sequences per backbone were designed using ProteinMPNN at a temperature of 0.1. Since Proteína is a Cα-only generative model, we used Cα conditioning (ca_only/v_48_020), whereas for RFdiffusion we used full backbone conditioning (vanilla_model_weights/v_48_020). We used the inference_ucond_400m_tri configuration for generating samples with Proteína and the default monomer configuration for RFdiffusion. Each sequence was refolded once using Boltz-1 with zero recycles, and 200 diffusion steps, and no MSA. A design was considered successful if it achieved a Boltz-1 predicted template modeling score (pTM) > 0.7 and a Cα RMSD < 2 Å between the refolded structure and the original generated backbone.

To ensure broad structural coverage, we clustered all designable backbones using Foldseek. This clustering is essential, as naive sampling strategies based on confidence scores alone result in under-representation of coils and structures with high β-sheet content (**Figure 2**)—a known issue in protein design, where non-helical structures typically score worse during design campaigns^22^. For curating our final experimental set, we combined all samples during clustering to ensure broad coverage of structural space, yielding 2,744 clusters.

For our final set of 384 designs, we sampled clusters with probability proportional to cluster size, preserving the natural output distribution of the generative models to assess their design space as-is. We alternated between Proteína and RFdiffusion, randomly selecting one design of the target model type from each sampled cluster; if a cluster contained no designs from the target model, we resampled a new cluster. For the comparison with the PDB (**Figure 2B**), deposited structures were clustered at 70% sequence identity level using MMseqs2^39^.

To compute metrics to correlate in-silico metrics to experimental outcomes (**Figure S4 and S8**), we refolded the 384 experimentally characterized sequences using 100 diffusion samples per model and 0 recycles and 100 recycles using Boltz-1, Boltz-2^40^, and AlphaFold3. For all 100 predictions, per-residue pLDDT scores were extracted and averaged across the structure. The pLDDT were computed both as whole-structure averages (HMQC correlations) and per-residue average (relaxation correlations). The RMSF (root-mean-square fluctuation) was computed by first aligning all 100 predicted models to a randomly chosen reference sample using Cα atoms. Per-residue RMSF values were then calculated as the standard deviation of atomic positions across the ensemble. The RMSF was computed for each atom in each residue in a structure and averaged over the residue. Mean (HMQC correlations) and per-residue (relaxation correlations) RMSF values were extracted.

Secondary structure fractions (helix, sheet, coil) were computed using DSSP^41^ on the predicted structures. The fraction of each secondary structure type was calculated as the number of residues assigned to that class divided by the total sequence length. Cα-RMSD between the designed backbone and refolded structure was computed after optimal superposition. For these metrics, we used only the original Boltz-1 refolded model, as this metric is primarily important for filtering designs that do not fold to the designed topology.

### Protein expression and purification

High-throughput cloning, expression and purification were adapted from Qian et al.^25^ with modifications for isotopic labelling necessary for NMR characterization. Designed amino acid sequences were reverse-translated and codon-optimized for expression in *E. coli* with DNA Chisel^42^, flanked with cloning adapters and ordered as synthetic DNA from Genscript. Gene fragments were assembled into a custom background-suppressing entry vector (Addgene plasmid #191551) using Golden Gate Assembly to append a C-terminal SNAC-tag followed by a His_6_-tag, and putting the designs under the control of a T7 promoter. Cloning reactions were performed in 1 µL volumes using BsaI-HFv2 (1.2 units, NEB #R3733), T4 ligase (40 units, NEB #M0202L), and gene fragments at a 2:1 molar ratio of insert to vector (4 fmol) in T4 ligase buffer. Reaction mixtures were prepared in 384-well source plates and transferred to a 96-well PCR plates (Eppendorf #0030129776) using an Echo 525 Acoustic Liquid Handler (Beckman Coulter). After assembly, reactions were incubated at 37°C for 20-60 minutes, followed by 5 min at 60°C.

Assembled plasmids were transformed into lab-made chemically competent BL21(DE3) *E. coli* cells (NEB #C2527) obtained with an adapted protocol from Yang et al.^43^ To each 1 µL Golden Gate reaction, 8 µL of ice-cold KCM (500 mM KCl, 250 mM MgCl_2_, 150 mM CaCl_2_) and 8 µL of ice-cold competent cells were added, followed by incubation on ice for 30 minutes. Heat shock was performed at 42°C for 10-15 seconds, followed by 5 minutes on ice. Cells were recovered in 100 µL SOC medium and incubated at 37°C for 1 hour with shaking at 1,000 rpm before proceeding directly to culture inoculation. We note that the lack of colony picking can in certain cases lead to polyclonal mixtures due to gene synthesis error. We therefore Sanger sequenced 48 samples. Based on a visual assessment of the sequencing chromatograms, 16% of samples indicated some level of polyclonality based on trace superposition at some locations.

For isotopically labeled protein expression, 25 µL of transformed cells were inoculated into 1 mL of LB supplemented with 100 µg/mL kanamycin in a 96-well deep-well plate (Fisherbrand #11391555) and grown at 37°C with shaking at 1,000 rpm for 12 h. From this starter plate, 25 µL of overnight culture was transferred into each well of eight replicate 96-well deep-well plates containing 1 mL per well of [^15^N]-labeled autoinduction medium. Composition of the autoinduction medium was adapted from Studier^44^ to be compatible with isotopic labeling. The medium contained 25 mM Na_2_HPO_4_, 25 mM KH_2_PO_4_, 5 mM Na_2_SO_4_, 2 mM MgSO_4_, 48 mM ^15^NH_4_Cl (2.6 g/L) as the sole nitrogen source, and an autoinduction sugar mix of 0.5% (v/v) glycerol, 0.05% (w/v) glucose, and 0.2% (w/v) lactose. The medium was supplemented with 0.2× trace metals (5 µM FeSO_4_, 0.5 µM ZnCl_2_, 0.5 µM Na_2_MoO_4_, 0.25 µM CaCl_2_, 0.21 µM H_3_BO_3_, 0.04 µM MnCl_2_, 0.02 µM CoCl_2_, 0.005 µM CuCl_2_) and 1× vitamin mix (1.5 mM NaCl, 11 µM i-inositol, 8.2 µM nicotinamide, 7.2 µM choline chloride, 4.9 µM pyridoxal HCl, 4.1 µM biotin, 3.0 µM thiamine HCl, 2.3 µM folic acid, 2.1 µM calcium pantothenate, 0.27 µM riboflavin) and 100 µg/ml Kanamycin. Plates were sealed with gas-permeable Breathe-Easiear sealing (Diversified Biotech #Z763624) and incubated at 37°C for 24 hours with shaking at 1,000 rpm.

Cultures were harvested by centrifugation at 4,000 × g for 5 minutes and pellets were lysed with BugBuster (Millipore #70921) supplemented with lysozyme (0.1 mg/mL), DNase I (0.01 mg/mL), and PMSF (1 mM) at 100 µL lysis buffer per 1 mL culture volume-equivalent. Lysis proceeded at 37°C for 15 minutes with shaking at 1,000 rpm until pellets were fully resuspended. Lysates were clarified by centrifugation at 4,000 × g for 15 minutes.

Protein purification was performed using Ni-NTA HisPur resin (Thermo Scientific #88222) in 96-well fritted plates (25 µm frit, Agilent #200953-100) using a vacuum manifold. 50 µL of resin bed volume were added to each well. The resin was equilibrated with three times 500 µL wash buffer (20 mM Tris-HCl pH 8.0, 300 mM NaCl, 25 mM imidazole). Clarified lysate was applied to equilibrated resin and incubated. Following binding, the resin was washed three times with 500 µL wash buffer. Excess buffer was removed by centrifugation at 1,000 × g for 1 minute. Bound proteins were eluted with 150 µL elution buffer (20 mM Tris-HCl pH 8.0, 300 mM NaCl, 500 mM imidazole) by centrifugation at 1,000 × g for 1 minute into 96-well collection plates.

Further purification was achieved by size exclusion chromatography on an Agilent 1260 Infinity II HPLC system equipped with a Superdex 75 Increase 5/150 column (Cytiva #29148722) equilibrated in SEC buffer (50 mM sodium phosphate, pH 6.8). Samples were filtered through 0.22 µm filter plates prior to injection. 100 µL of each sample were injected at a flow rate of 0.65 mL/min. Elutions were monitored at 280 nm and fractions corresponding to monomeric protein peaks were identified based on a calibration curve. Appropriate fractions were pooled automatically using an Opentrons Flex liquid handling robot before downstream NMR analysis. The protein samples were adjusted to 475 μL, the D_2_O content was adjusted to 5% and the DSS-d_4_ concentration to 100 μM. All samples were transferred into standard 5 mm tubes for subsequent NMR analysis.

The 9 proteins characterized in detail were manually re-purified from an expression volume of 100 mL, with ^13^C-glucose added during expression, and using IPTG for induction. Affinity purification was performed on a 5 mL HisTrap column (Cytiva) with collection of 0.5 mL fractions. The protein-containing fractions were pooled and concentrated to 0.5 mL using centrifugation with Amicon4 3k MWCO (Merck Millipore), and then purified using a Superdex 75 10/300 column (Cytiva #29148721), collecting 0.25 mL fractions.

### NMR spectroscopy

Spectra were automatically acquired with IconNMR in Topspin 3.7.0, utilizing a 24-sample storage Bruker SampleCase. All samples were temperature-equilibrated for 1 minute after being placed in the NMR magnet, followed by automated sample setup. Automated calibration of the proton 90° pulse (p1) and the water frequency offset (o1) was performed using the Bruker au macro “au_zg”. All experiments were carried out at 25°C on a Bruker AVANCE III HD spectrometer operating at 600.13 MHz equipped with a cryogenic triple-resonance probe. For the standard 2D [^15^N, _1_H]-SOFAST HMQC measured on all 379 samples, the ^1^H carrier was centered on the water resonance, the ^15^N carrier at 118 ppm. The interscan delay was set to 0.2 s. In the direct dimension, 512 complex points were recorded in an acquisition time of 53 ms, multiplied with a 90°-shifted sine square bell, zero-filled to 1024 points and Fourier transformed. In the indirect dimension, 256 complex points were measured with a maximal evolution time of 124 ms, multiplied with a 90°-shifted sine bell, zero-filled to 512 points and Fourier transformed. Solvent filter and polynomial baseline correction were applied in the direct dimension.

For assignment and structure determination, the following NMR experiments from the NMRlib^45^ were recorded: 2D [_13_C, _1_H]-HSQC, 3D BEST-HNCACB, 3D [^1^H,_1_H]-NOESY-^15^N-HSQC, 3D [^1^H, ^1^H]-NOESY-_13_C-HSQC, 3D HccoNH-TOCSY, 3D hCcoNH-TOCSY, ^15^N TRACT. For relaxation, ^15^N *T*_1_-HSQC, ^15^N *T*_2_-HSQC, and ^15^N{^1^H}-NOE. Additional 4D TROSY APSY-HNCOCA (oaNH) and 1D Proton NMR spectra with excitation sculpting were recorded using pulse programs provided by the Bruker standard library.

### Structure calculation

Structure calculation was performed with CYANA 3.98.15^46^. All spectra were processed and analyzed with NMRPipe^47^ and ccpNMR^48^ version 3. Peak picking for the 3D [^1^H,^1^H]-NOESY-^15^N-HSQC and the 3D [^1^H,^1^H]-NOESY-^13^C-HSQC was performed using ARTINA^49^ peak picking on the NMRtist.org webservice, yielding a total of 2091 experimental NOEs. 184 dihedral constraints were derived by TALOS-N^50^ from the experimentally determined C_α_, C_β_, N and HN chemical shifts. A total of 200 structures were calculated, and the ensemble of 20 lowest energy structures was selected. Ramachandran statistics of this ensemble showed 90.8% of residues in most favored regions, 9.2% in allowed regions, 0% in generously allowed regions, and 0% in disallowed regions.

### Quality estimation

Spectral quality was categorized using a combined score. Peaks were picked using the peak picker of the NMRglue^51^ Python package, yielding a total number of peaks N_P_. All spectra with N_P_ ≤ 2 were classified as “no signal”. The number of expected peaks N_E_ was calculated as N_E_ = L – 1 – n(P) + 2n(N) + 2n(Q) + n(W), where L is the length of the sequence, and n(P), n(N), n(Q), and n(W) are the numbers of proline, asparagine, glutamine, and tryptophan, respectively. The peak miss score was calculated as p_M_ = |10 * (N_E_ – N_P_) /N_E_|. The noise level of the spectrum was defined as the median absolute deviation of the full spectrum. The score for the median signal-to-noise ratio of all peaks (MI) was calculated as p_I_ = (35 – MI) /10, where 0 < p_I_ < 3.5. A total score p = p_I_ + p_M_ < 3 was considered “Very good”, 3 ≤ p < 6 “Good”, and p ≥6 “Poor”.

### Secondary structure subgrouping

Classification of the designed proteins based on their structure was done by automated calculation of backbone angles from the designed structure. Residues with ϕ between –160° and –30° and ψ between –100° and 50° were categorized as α-helix, those with ϕ between – 180° and –40° and ψ either between 70° and 180° or –180° and –120° as β-sheet. The percentages of helix and sheet in the structure gave rise to the five categories “α-helix”, “mainly α”, “mixed”, “mainly β” and “β-sheet” (SI Table 1).

### HMQC spectral clustering and dendrogram construction

To quantify spectral similarity across the designed protein library, we computed pairwise distances between two-dimensional ^1^H-^15^N HMQC peak lists using the symmetric Chamfer distance. The Chamfer distance is defined as:

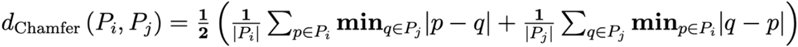

, where *P*_*i*_ and *P*_*j*_ are the sets of picked peaks (each a 2D coordinate in ^1^H and ^15^N chemical shift space) for spectra *i* and *j*, respectively. Prior to distance computation, ^1^H and ^15^N chemical shifts were independently z-score normalized across all samples to place both dimensions on a comparable scale (^1^H shifts span approximately 6–12 ppm; ^15^N shifts span approximately 90–140 ppm).

The resulting symmetric distance matrix (238 × 238) was converted to a condensed form and subjected to agglomerative hierarchical clustering using Ward’s minimum-variance linkage. The dendrogram was rendered with leaves ordered by the optimal leaf-ordering algorithm and annotated with a categorical color bar indicating predicted fold class. Fold class was assigned from DSSP secondary-structure strings computed on Boltz-predicted structures: designs with ≥40% helical content and <10% strand content were classified as “Mostly α”, and all others as “Mixed α/β.”

## Figures

All images of protein structures were created using UCSF ChimeraX^52^. All plots were created with the Python package Matplotlib^53^. For additional data analysis, we used the python packages biotite^54^ and scipy^55^.

