## Supplementary Information for "Large-scale exploration of protein space by automated NMR"

This file contains:

- Supplementary Tables S1–S2
- Supplementary Figures S1–S8

**Supplementary Table S1.** NMR and refinement statistics for p2 A4.

| | p2 A4 |
|---|---|
| NMR distance and dihedral constraints | 2275 |
| Distance constraints | 2091 |
| Total NOE | 2091 |
| Intra-residue | 409 |
| Inter-residue | 1682 |
| Sequential ($\|i - j\| = 1$) | 592 |
| Medium-range ($1 < \|i - j\| < 5$) | 469 |
| Long-range ($\|i - j\| >= 5$) | 621 |
| Intermolecular | 0 |
| Hydrogen bonds | 0 |
| Total dihedral angle restraints | 184 |
| $\phi$ | 92 |
| $\psi$ | 92 |
| Structure statistics | |
| Violations (mean and s.d.) | |
| Distance constraints (Å) | $0.008 \pm 0.03$ |
| Dihedral angle constraints (°) | $0.87 \pm 0.12$ |
| Max. dihedral angle violation (°) | $5.34 \pm 1.58$ |
| Max. distance constraint violation (Å) | $0.30 \pm 0.21$ |
| Deviations from idealized geometry* | |
| Average pairwise r.m.s. deviation** (Å) | |
| Heavy | $0.40 \pm 0.10$ |
| Backbone | $0.73 \pm 0.11$ |

* There are no covalent bond-length or bond-angle outliers.

** Calculated among 20 refined structures

**Supplementary Table S2.** Protein categories based on secondary structure

| | **α-helix** | **mainly α** | **mixed** | **mainly β** | **β-sheet** |
|---|---|---|---|---|---|
| **% sheet** | 0 | > 0–30 | > 30–60 | 40–79 | 66–82 |
| **% helix** | 77–95 | 50–91 | > 30–60 | > 0–30 | 0 |
| **count** | 141 | 134 | 76 | 29 | 4 |

**Supplementary Figures**

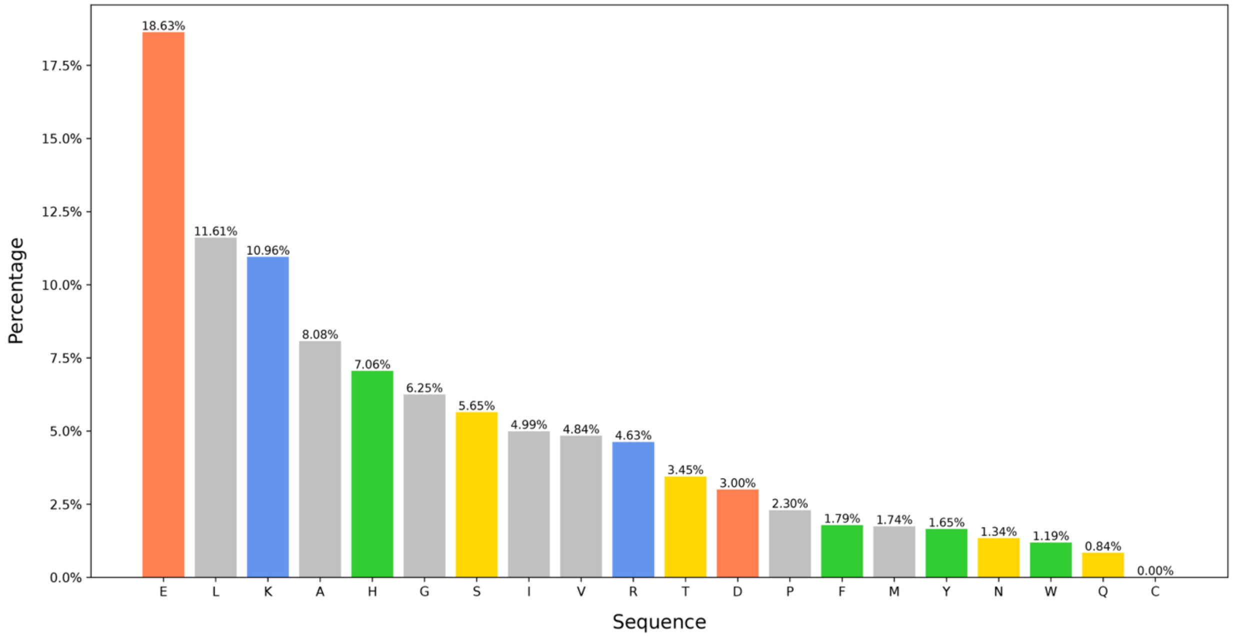


**Supplementary Figure S1.** Relative abundance of each of the 20 standard amino acids amongst all 384 designed proteins. Basic amino acids are shown in blue, acidic in red, aromatic in green, polar in yellow, and nonpolar in gray.

p1 A4

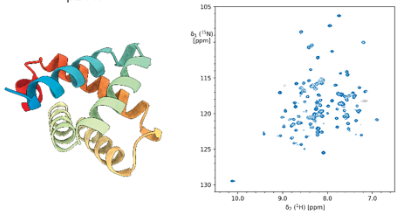

p1 A7

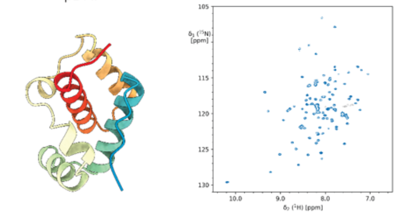

p1 A8

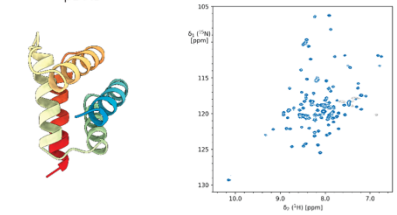

p1 A10

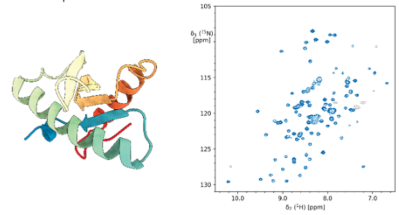

p1 A11

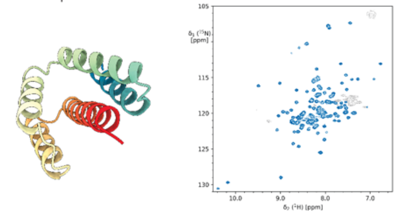

p1 A12

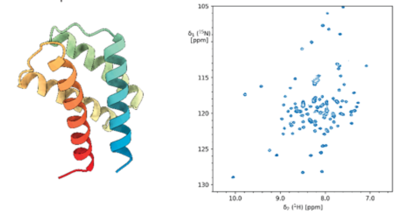

p1 B1

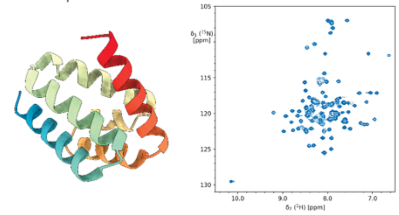

p1 B2

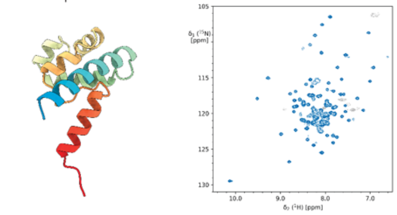

p1 B3

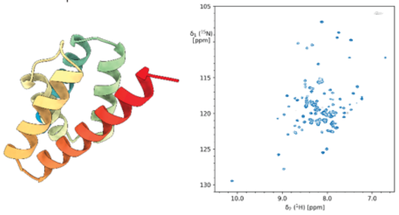

p1 B4

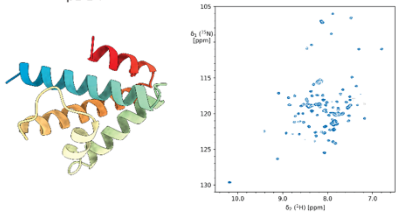

p1 B5

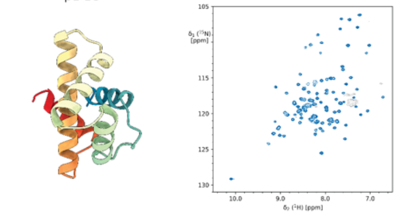

p1 B6

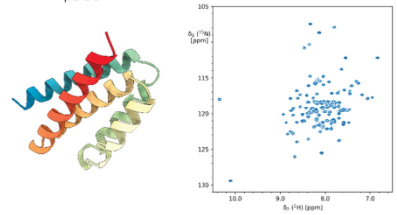

p1 B7

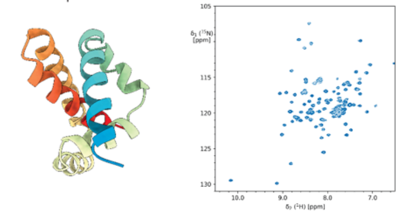

p1 B8

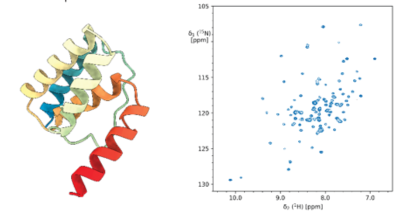

p1 B10

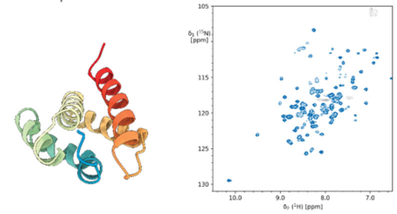

p1 B11

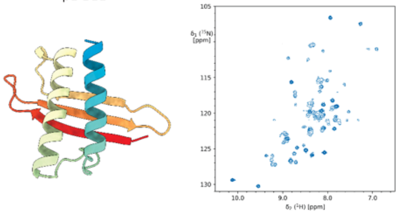

p1 B12

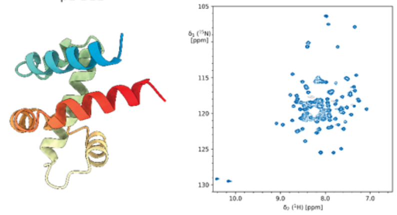

p1 C1

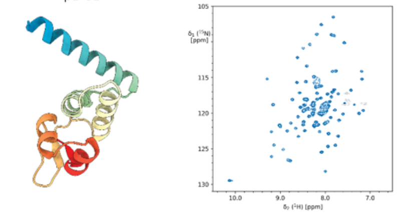

p1 C2

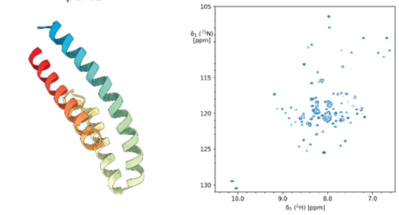

p1 C4

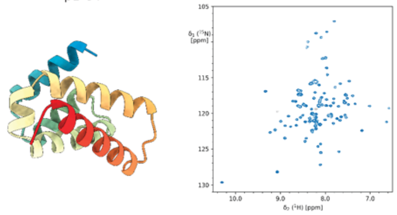

p1 C6

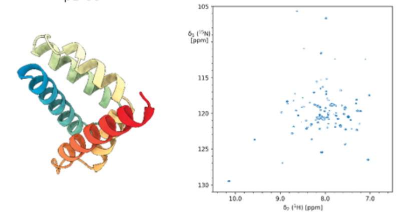

p1 C7

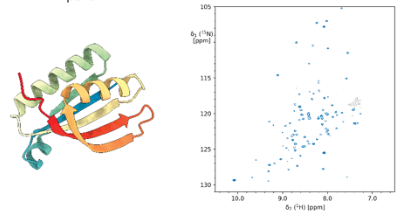

p1 C10

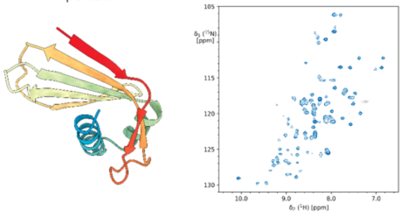

p1 C11

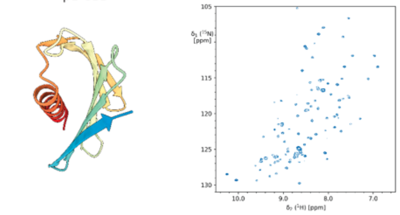

p1 D1

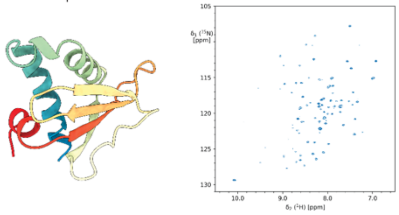

p1 D3

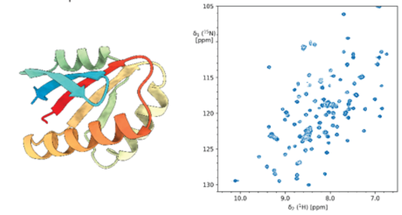

p1 D4

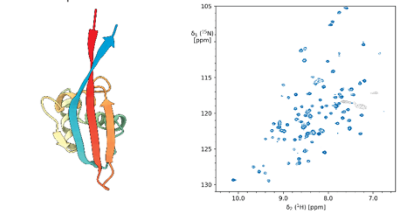

p1 D6

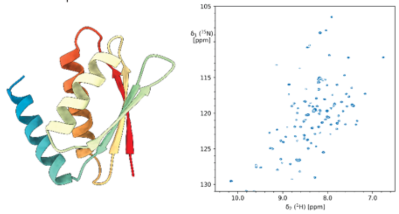

p1 D7

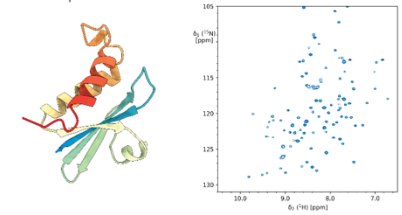

p1 D8

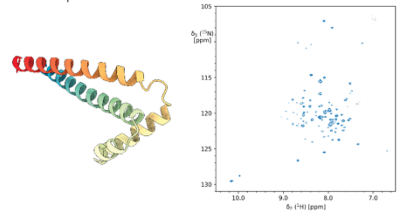

p1 D9

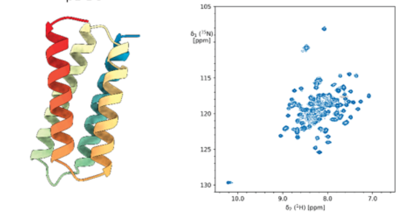

p1 D10

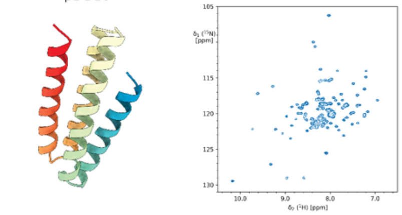

p1 D11

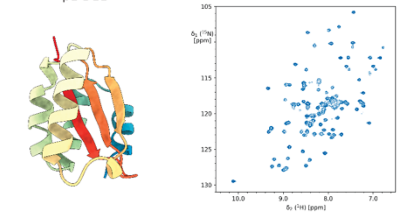

p1 D12

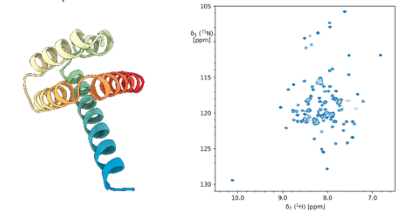

p1 E2

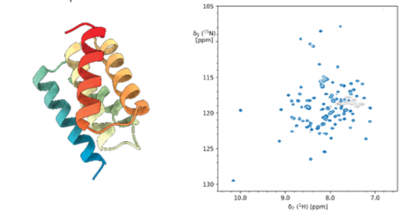

p1 E4

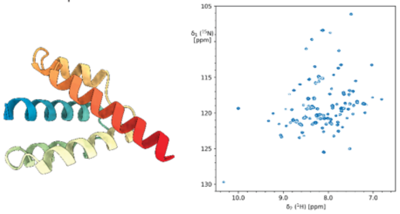

p1 E7

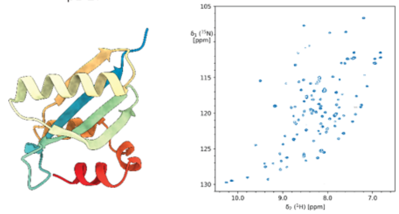

p1 E9

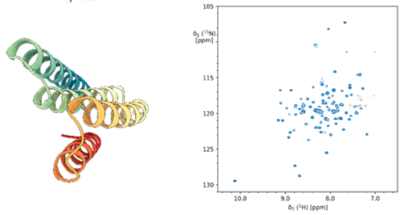

p1 E12

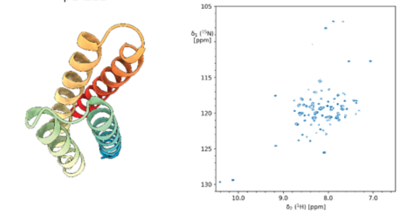

p1 F2

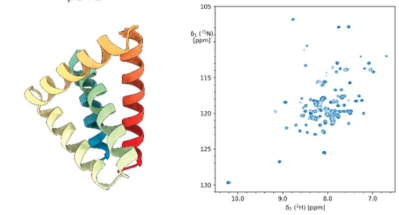

p1 F3

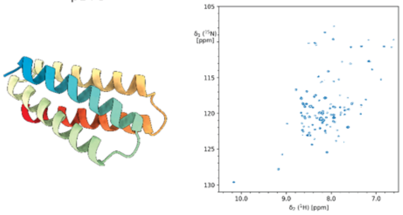

p1 F5

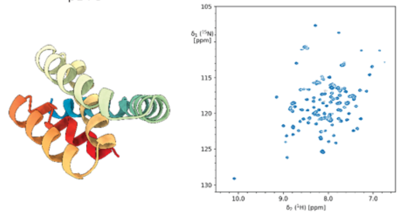

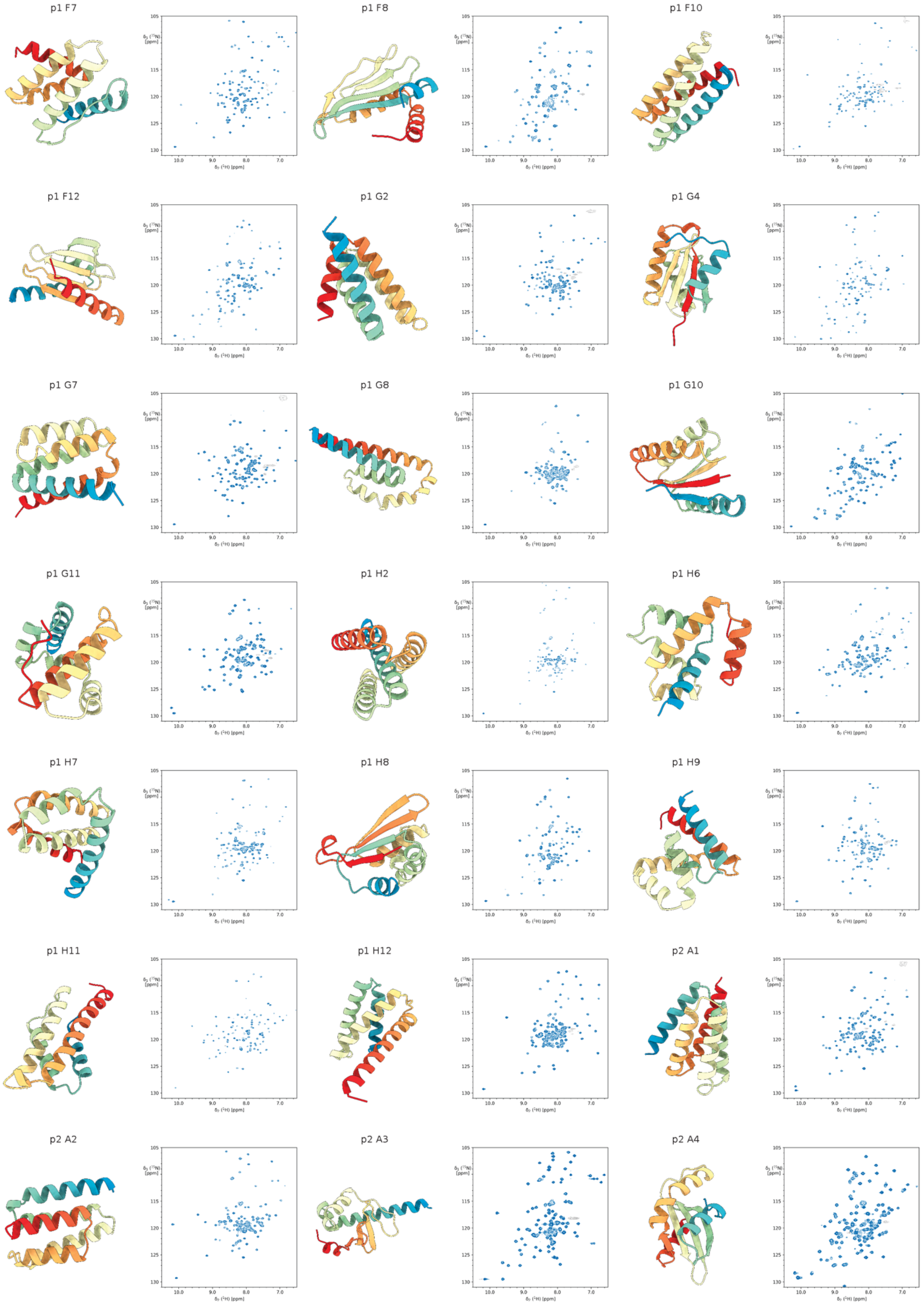
p1 F7
p1 F8
p1 F10
p1 F12
p1 G2
p1 G4
p1 G7
p1 G8
p1 G10
p1 G11
p1 H2
p1 H6
p1 H7
p1 H8
p1 H9
p1 H11
p1 H12
p2 A1
p2 A2
p2 A3
p2 A4

p2 A5

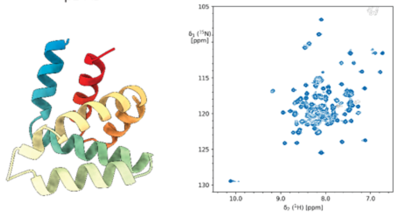

p2 A6

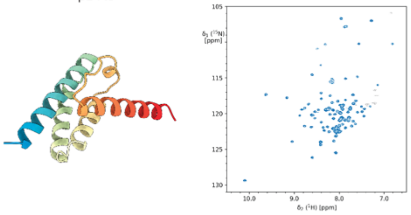

p2 A7

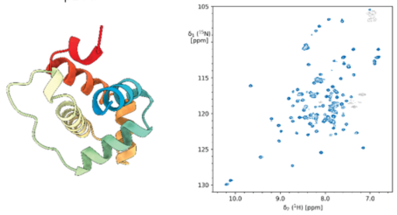

p2 A8

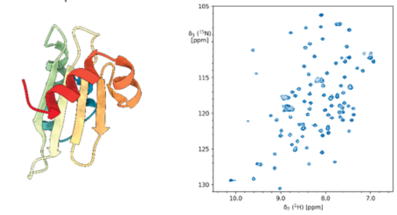

p2 A9

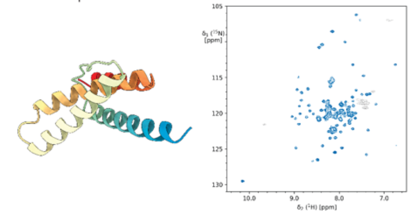

p2 A10

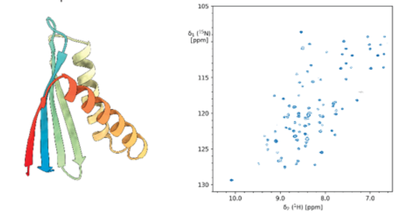

p2 A12

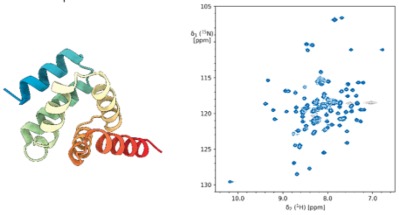

p2 B2

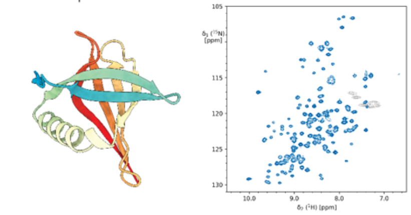

p2 B3

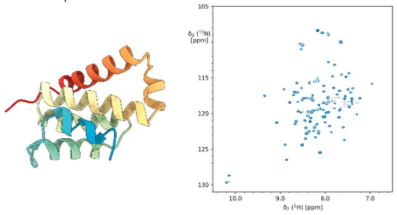

p2 B4

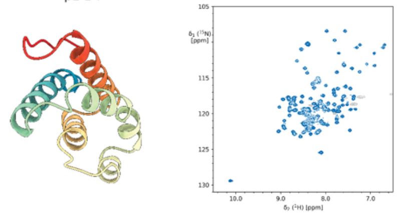

p2 B5

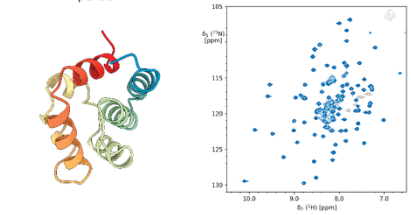

p2 B7

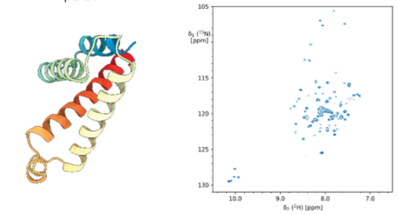

p2 B8

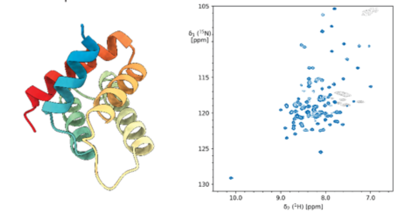

p2 B9

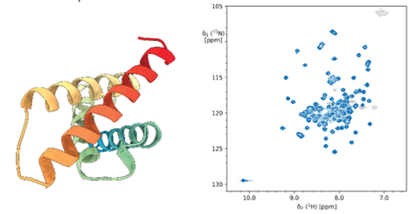

p2 B10

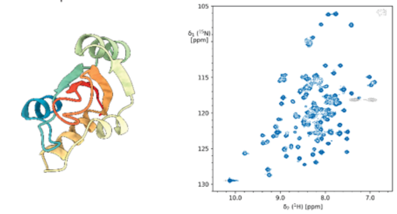

p2 B11

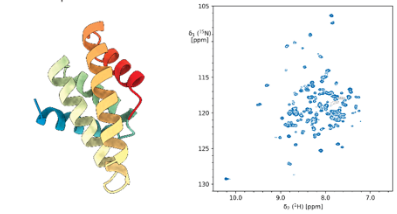

p2 B12

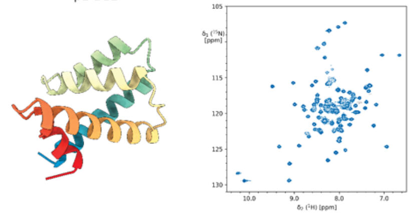

p2 C2

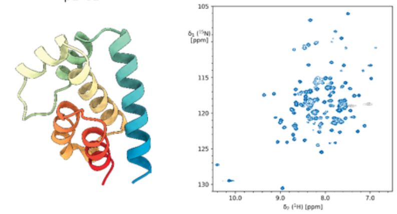

p2 C3

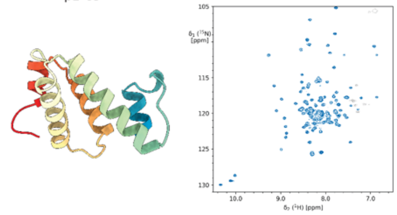

p2 C5

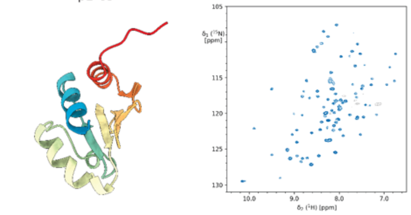

p2 C6

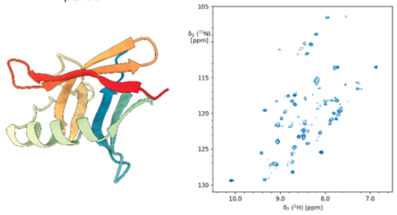

p2 C9

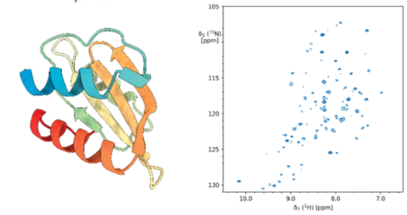

p2 C11

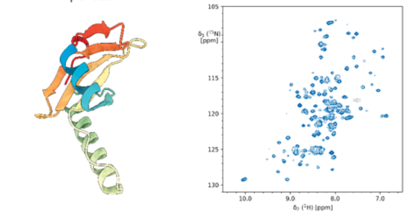

p2 D1

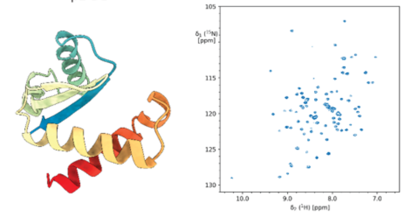

p2 D3

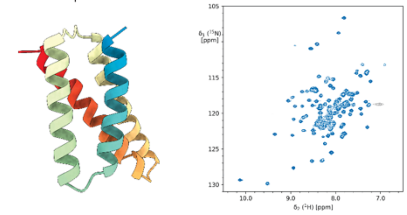

p2 D4

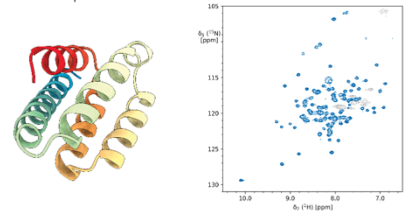

p2 D5

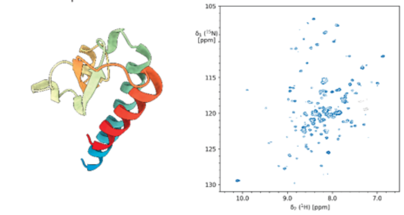

p2 D6

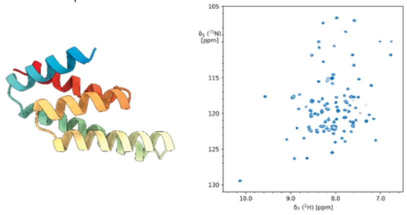

p2 D7

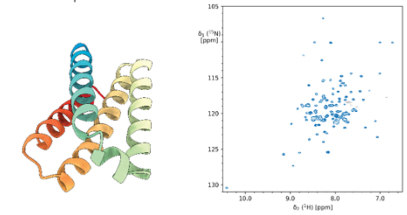

p2 D8

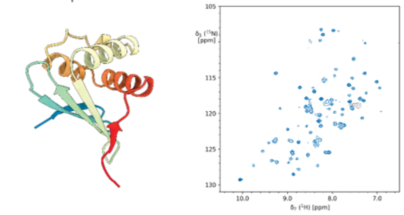

p2 D9

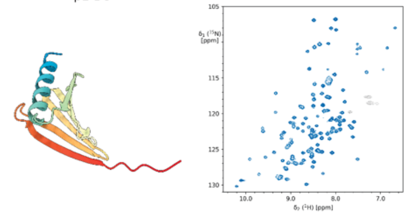

p2 D10

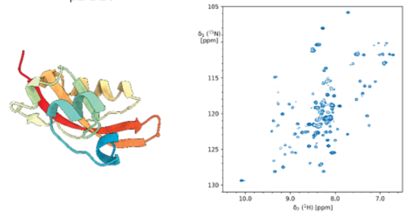

p2 D11

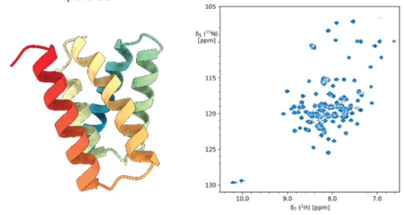

p2 D12

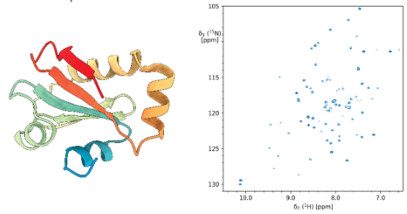

p2 E1

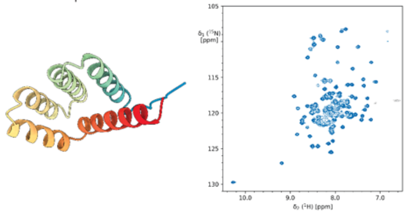

p2 E3

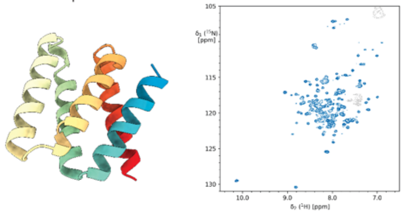

p2 E5

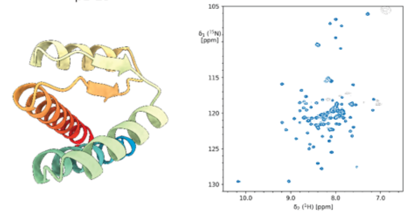

p2 E6

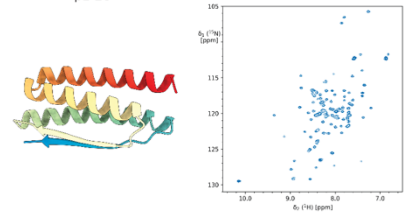

p2 E8

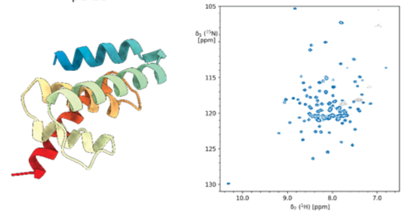

p2 E9

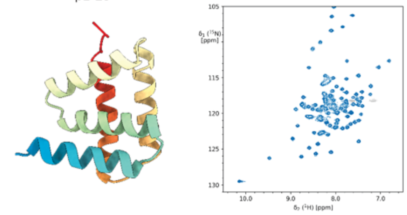

p2 E10

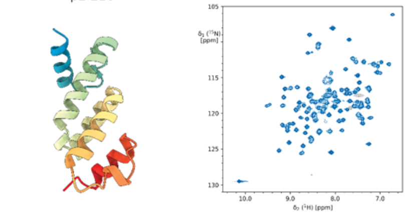

p2 E11

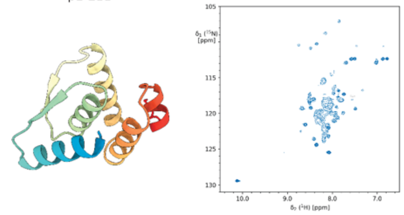

p2 E12

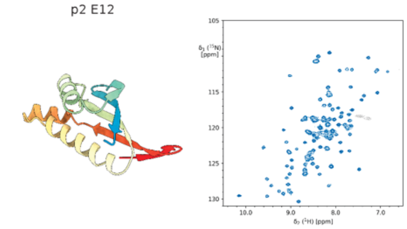

p2 F1

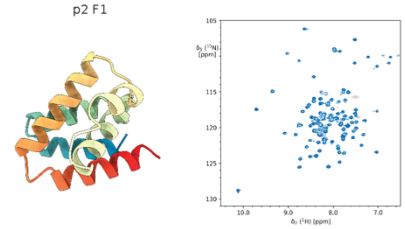

p2 F2

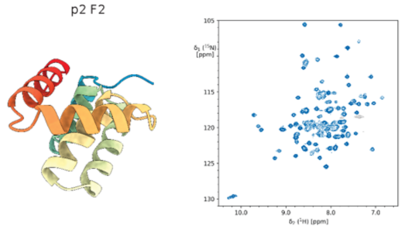

p2 F4

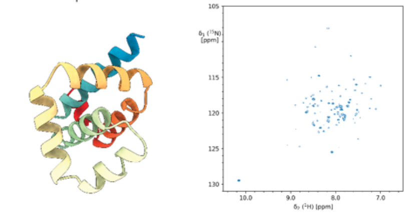

p2 F5

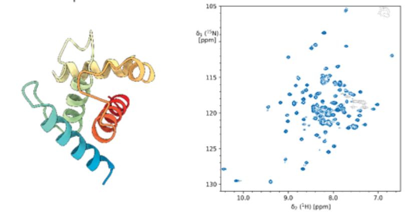

p2 F9

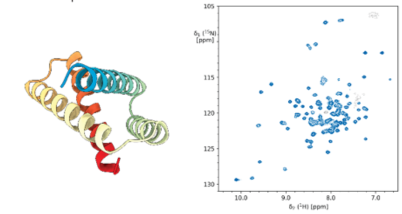

p2 F12

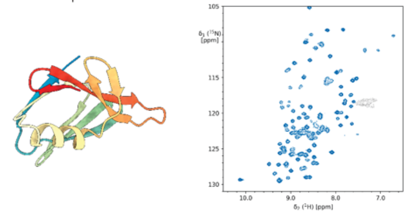

p2 G1

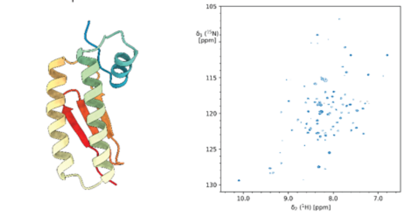

p2 G2

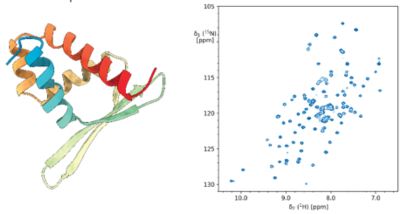

p2 G4

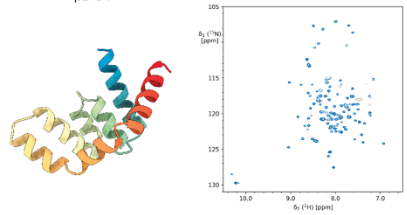

p2 G6

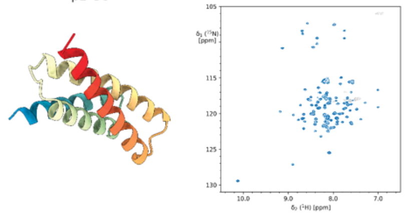

p2 G7

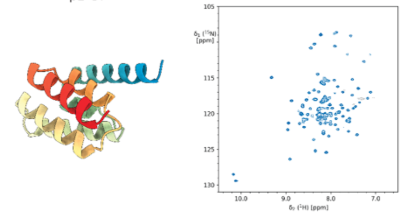

p2 G9

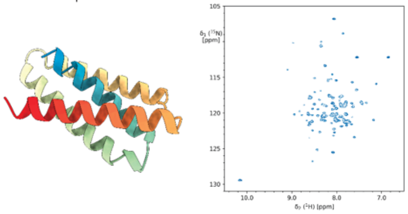

p2 G11

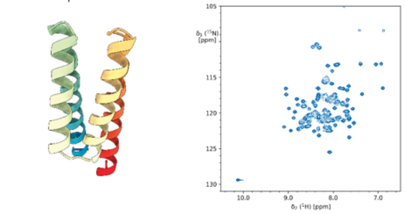

p2 G12

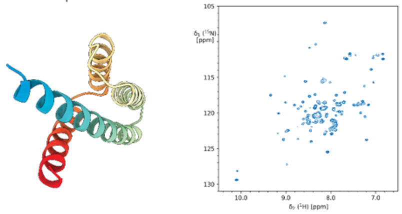

p2 H1

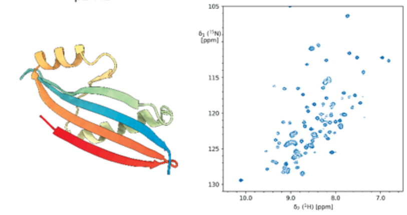

p2 H2

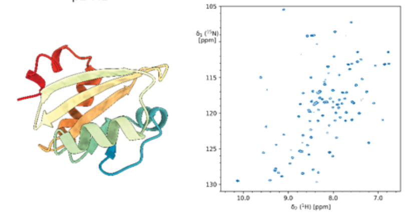

p2 H3

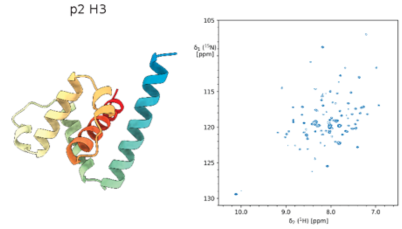

p2 H6

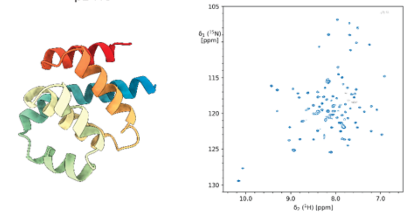

p2 H7

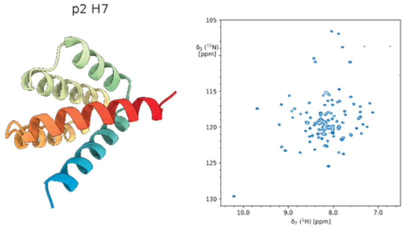

p2 H8

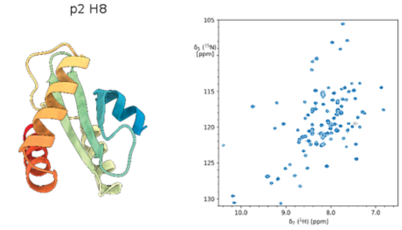

p2 H11

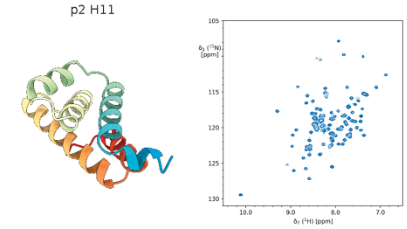

p2 H12

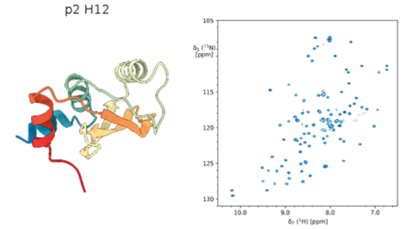

p3 A1

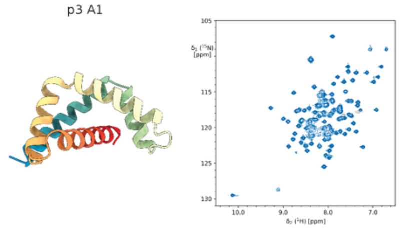

p3 A3

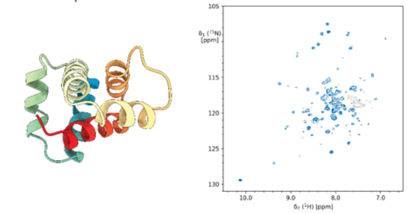

p3 A7

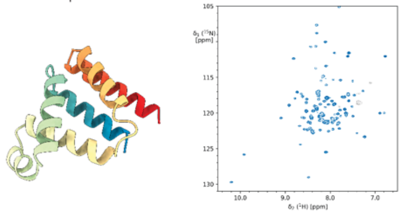

p3 A8

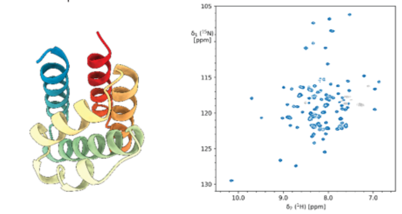

p3 A10

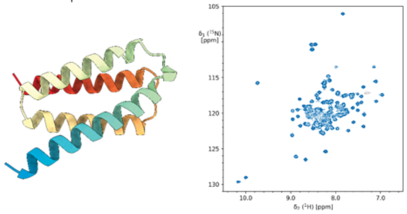

p3 A11

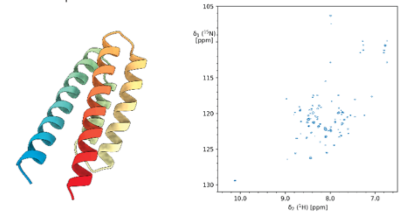

p3 A12

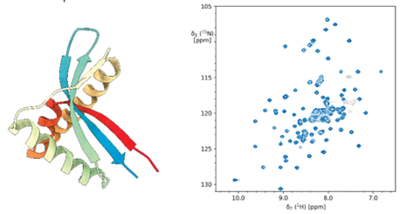

p3 B1

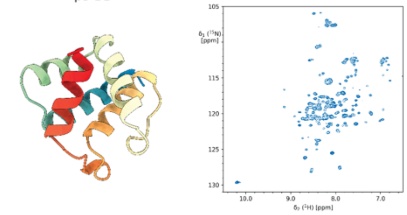

p3 B2

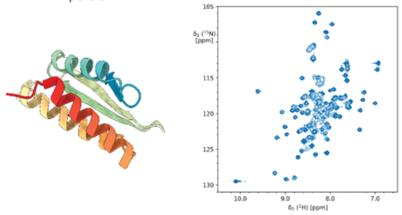

p3 B5

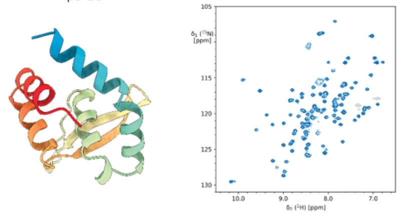

p3 B6

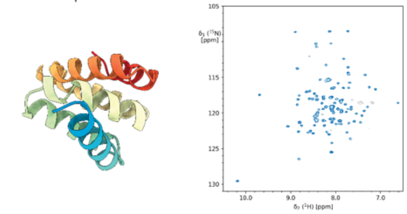

p3 B7

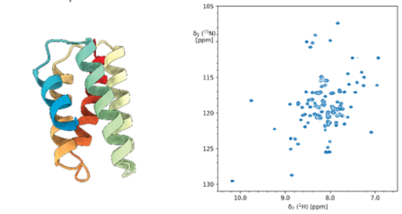

p3 B9

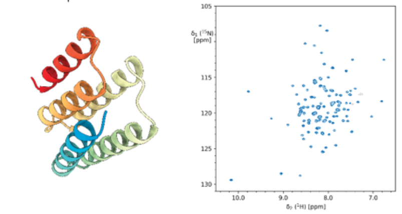

p3 B10

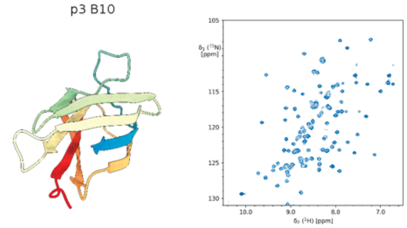

p3 B11

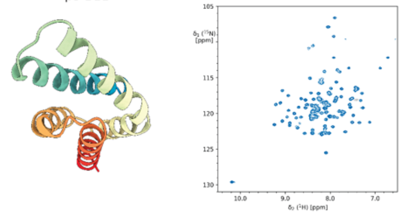

p3 B12

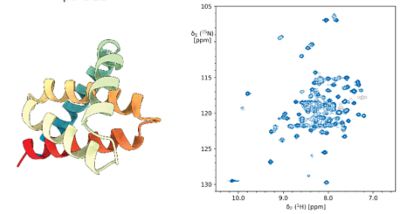

p3 C2

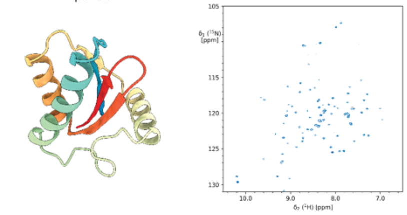

p3 C5

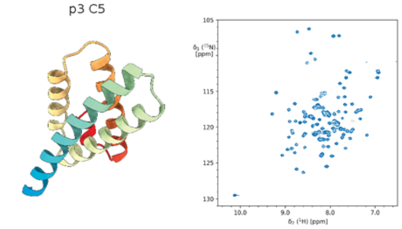

p3 C7

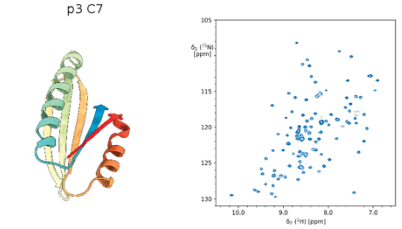

p3 C9

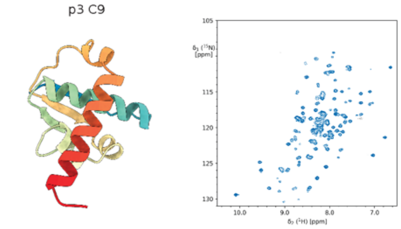

p3 C10

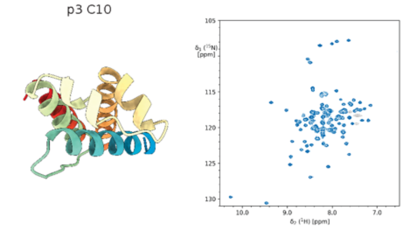

p3 C11

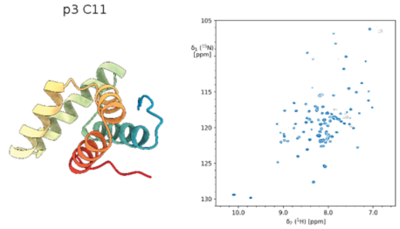

p3 D2

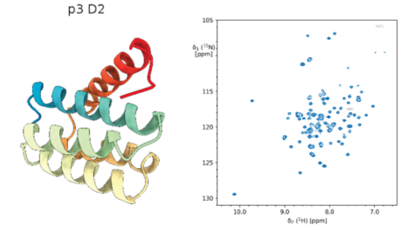

p3 D4

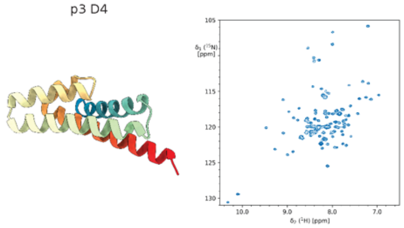

p3 D6

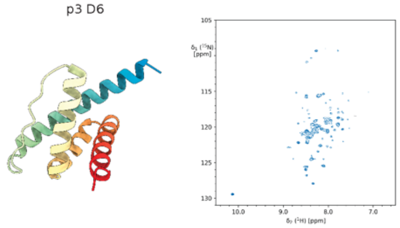

p3 D10

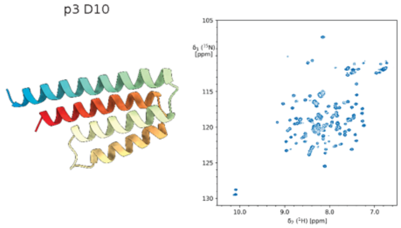

p3 D11

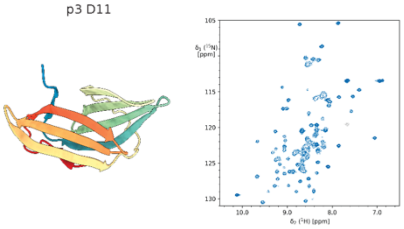

p3 D12

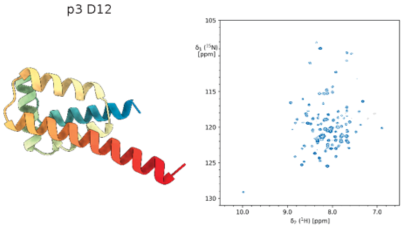

p3 E1

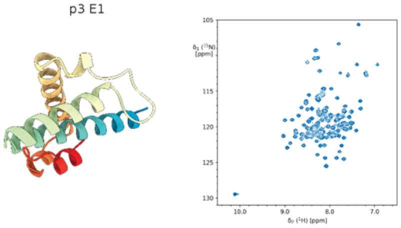

p3 E3

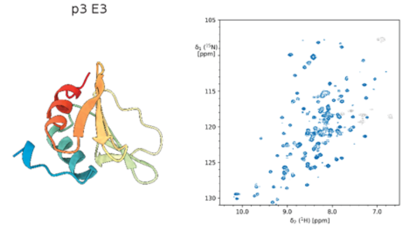

p3 E5

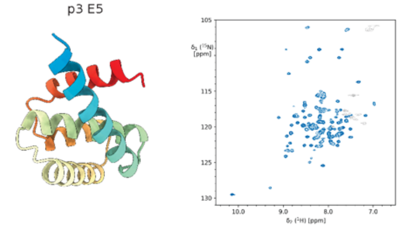

p3 E6

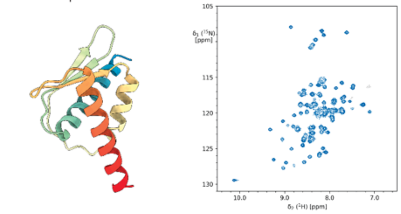

p3 E10

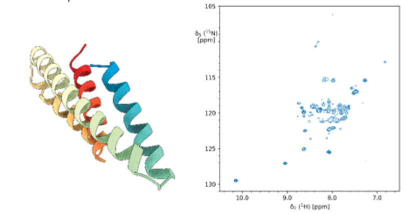

p3 E11

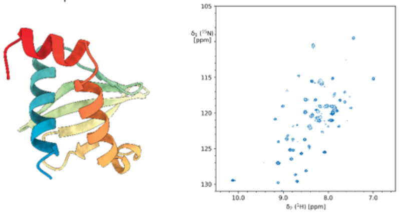

p3 E12

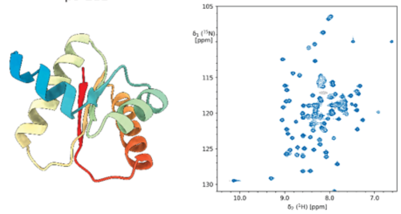

p3 F1

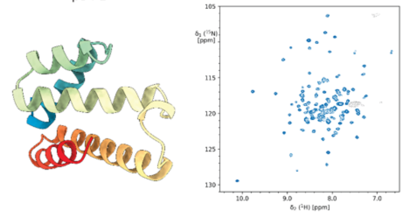

p3 F2

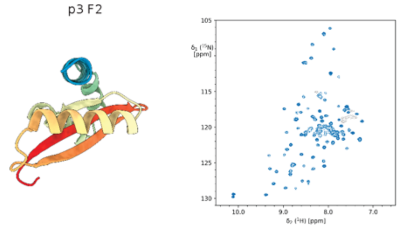

p3 F8

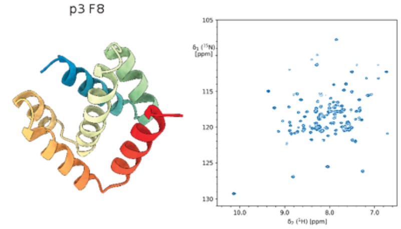

p3 F9

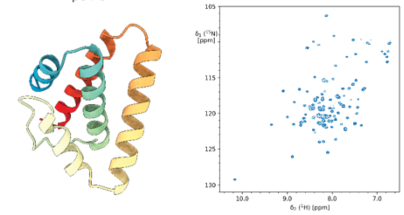

p3 F12

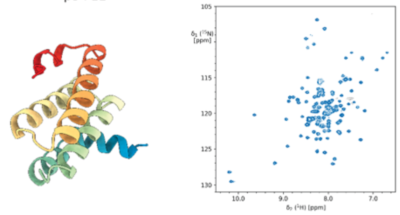

p3 G1

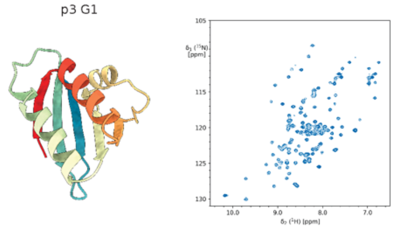

p3 G2

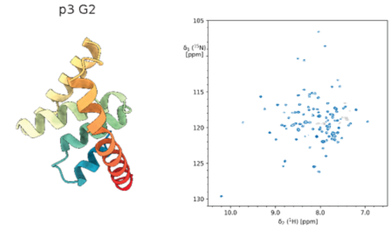

p3 G5

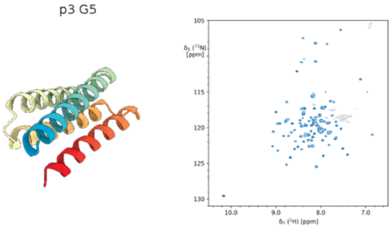

p3 G9

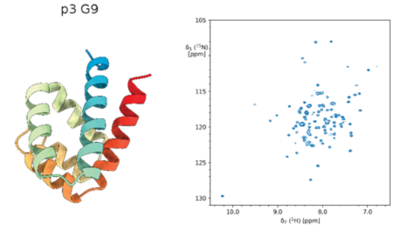

p3 G11

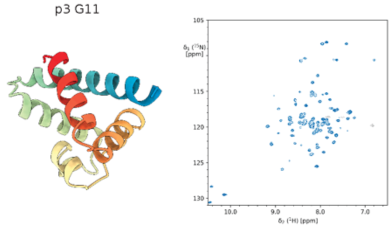

p3 G12

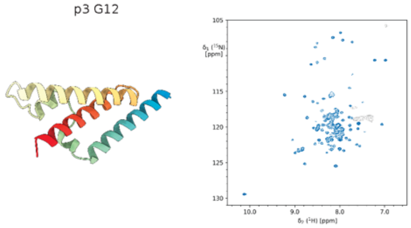

p3 H2

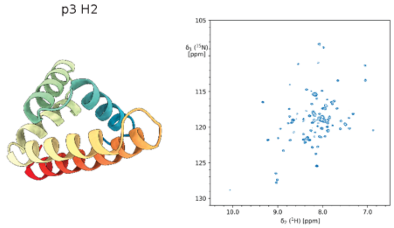

p3 H4

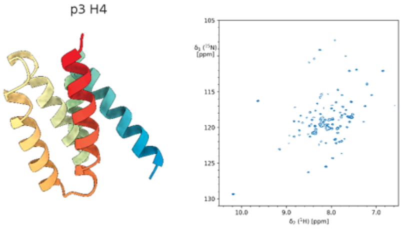

p3 H5

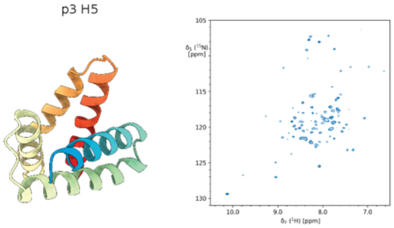

p3 H6

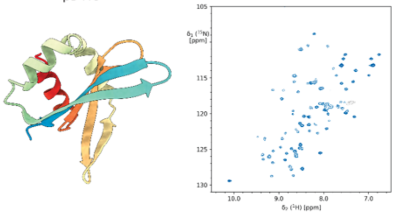

p3 H7

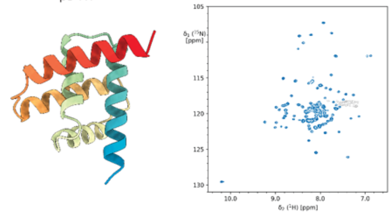

p3 H9

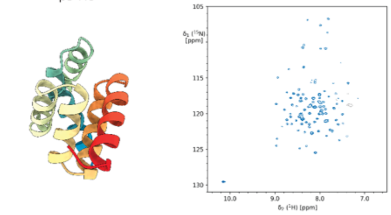

p3 H10

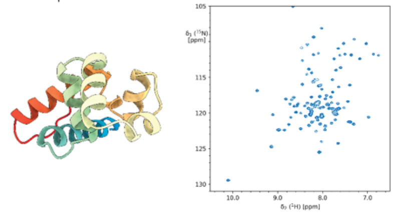

p4 A5

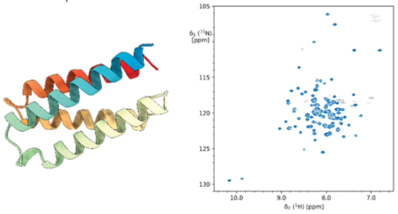

p4 A7

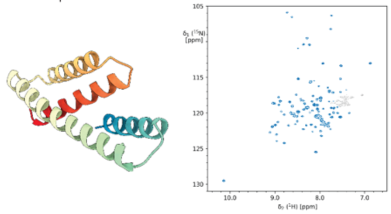

p4 A8

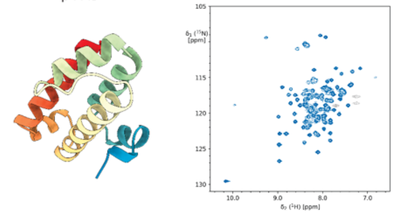

p4 A9

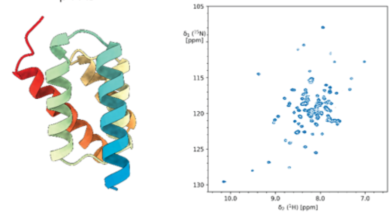

p4 B2

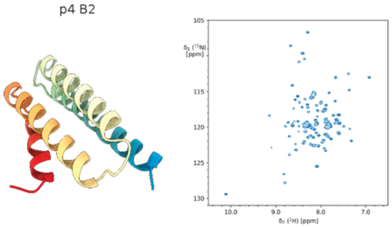

p4 B4

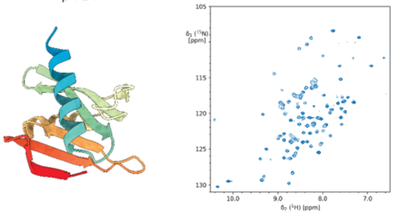

p4 B6

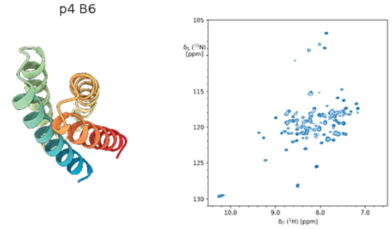

p4 B7

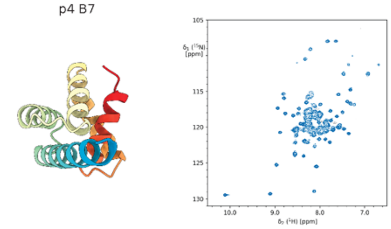

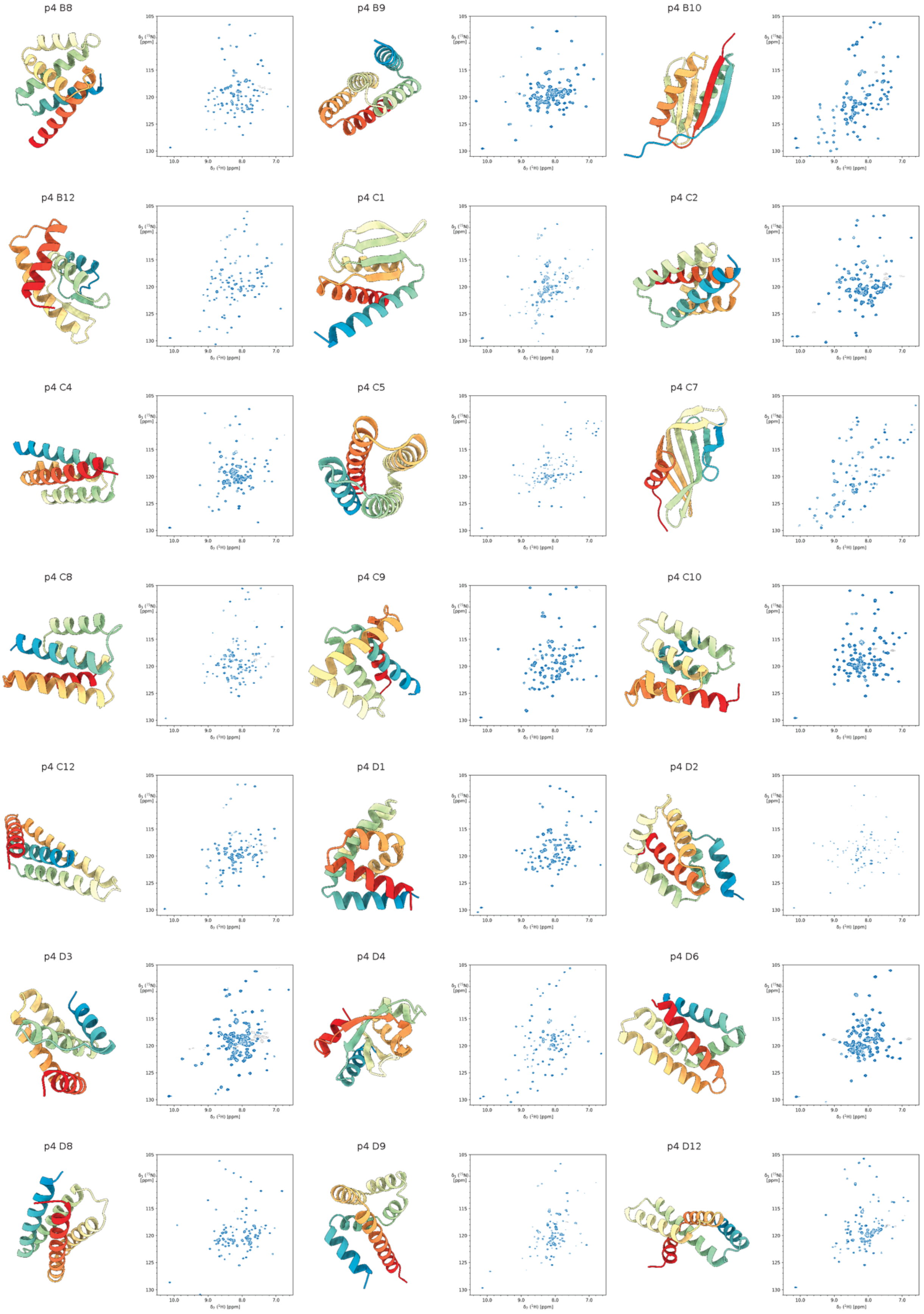
p4 B8
p4 B9
p4 B10
p4 B12
p4 C1
p4 C2
p4 C4
p4 C5
p4 C7
p4 C8
p4 C9
p4 C10
p4 C12
p4 D1
p4 D2
p4 D3
p4 D4
p4 D6
p4 D8
p4 D9
p4 D12

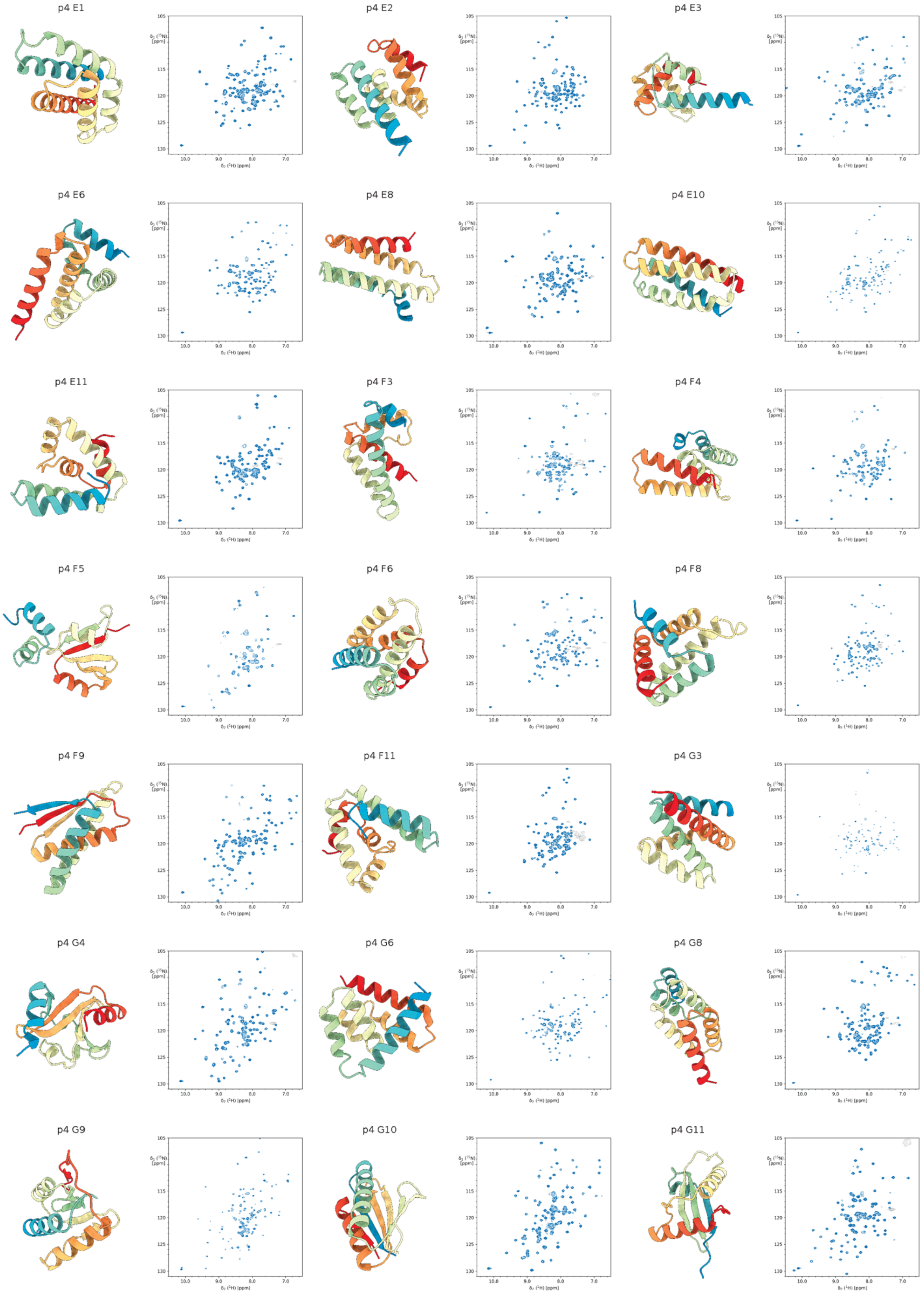
p4 E1
p4 E2
p4 E3
p4 E6
p4 E8
p4 E10
p4 E11
p4 F3
p4 F4
p4 F5
p4 F6
p4 F8
p4 F9
p4 F11
p4 G3
p4 G4
p4 G6
p4 G8
p4 G9
p4 G10
p4 G11

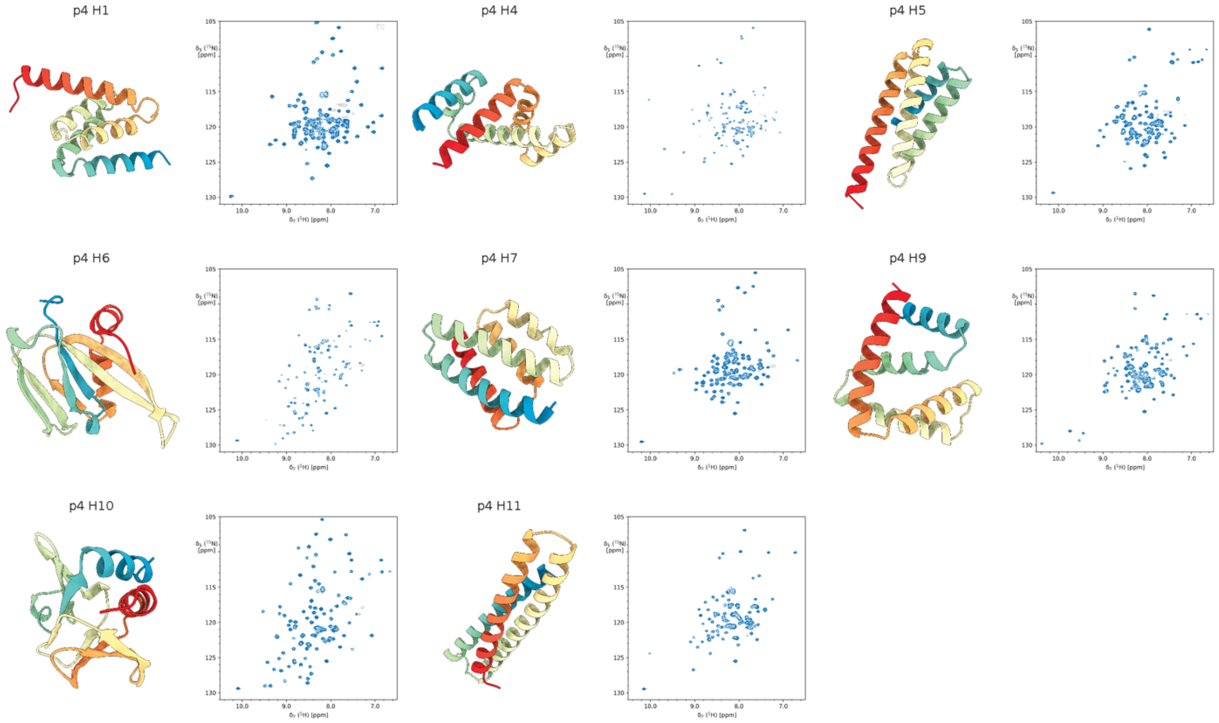


**Supplementary Figure S2.** All spectra and structures classified as good (“good” and “very good”).

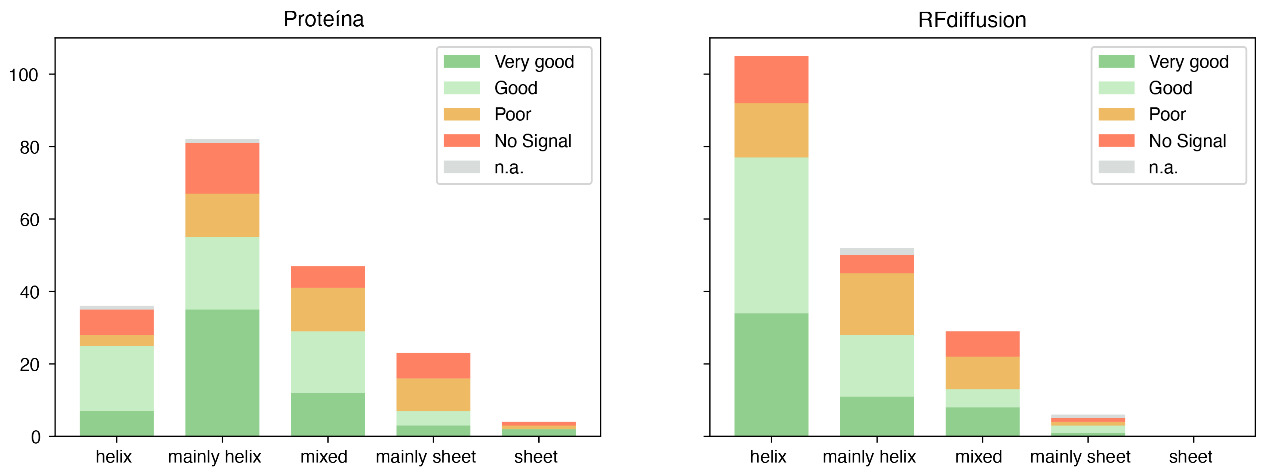


**Supplementary Figure S3.** Numbers and quality distribution of designed proteins per secondary structure subgroup for each design model.

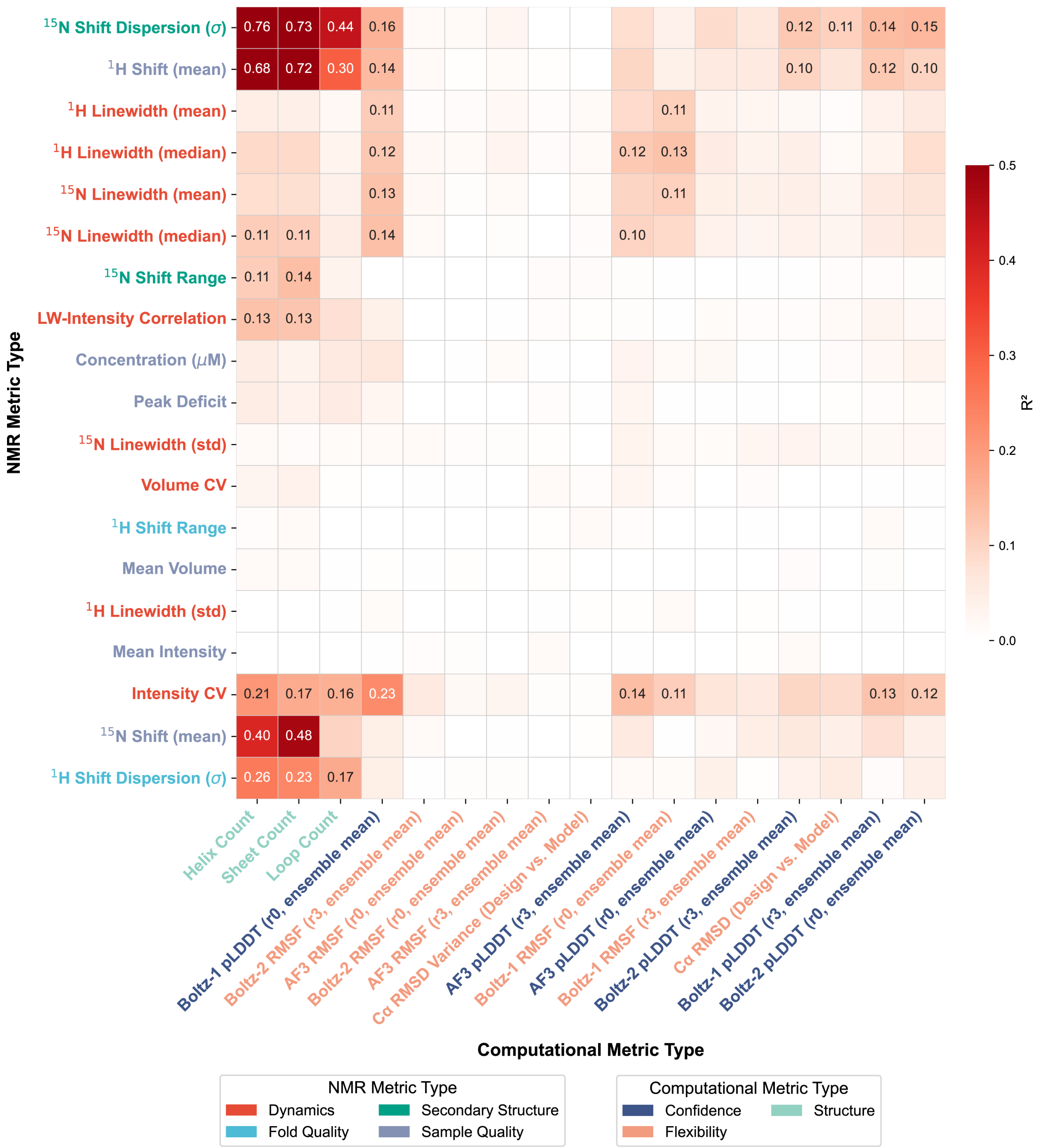


**Supplementary Figure S4.** Correlation between computational metrics and NMR spectrum-level experimental observables. Computational metrics are shown on the x-axis, color-coded by category. Experimental observables at the level of HMQC spectra are shown on the y-axis, color-coded by category. Coefficient of determination ($R^2$) are indicated as a colormap for each pair of computational-experimental metric. Coefficient greater than 0.1 are indicated explicitly as numerical values.

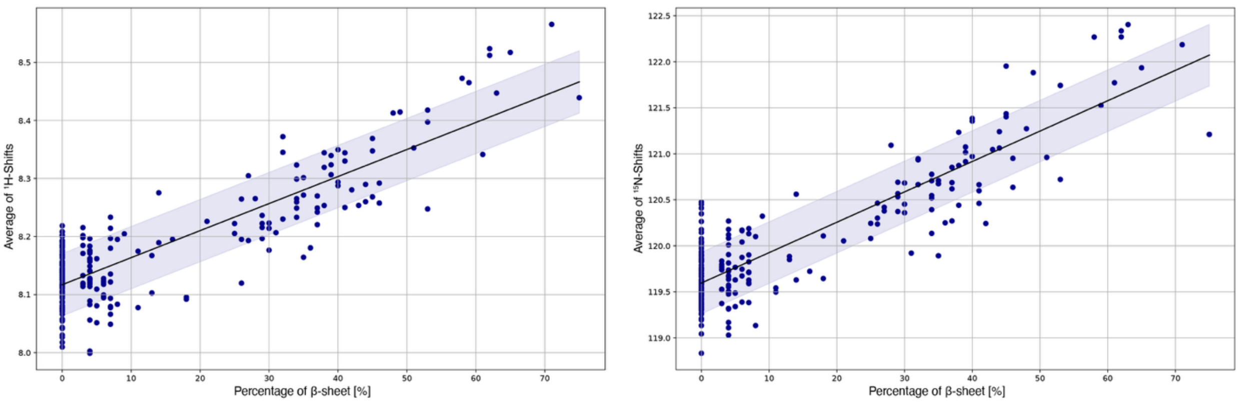


**Supplementary Figure S5.** Average chemical shifts as a function of the percentage of β-sheet, for the proteins classified "good" and "very good".

p2 A4

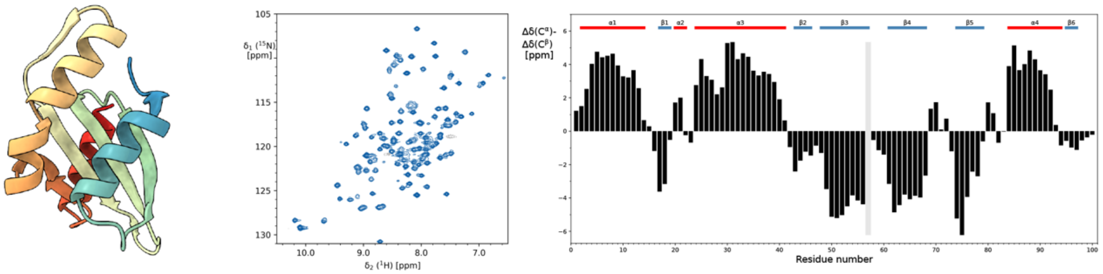


p2 E10

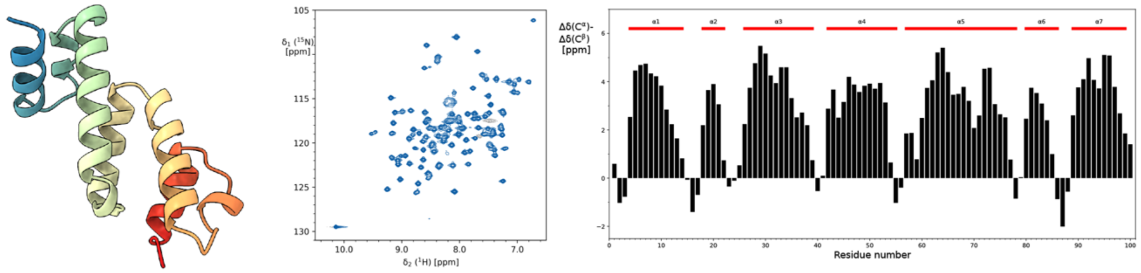


p3 A1

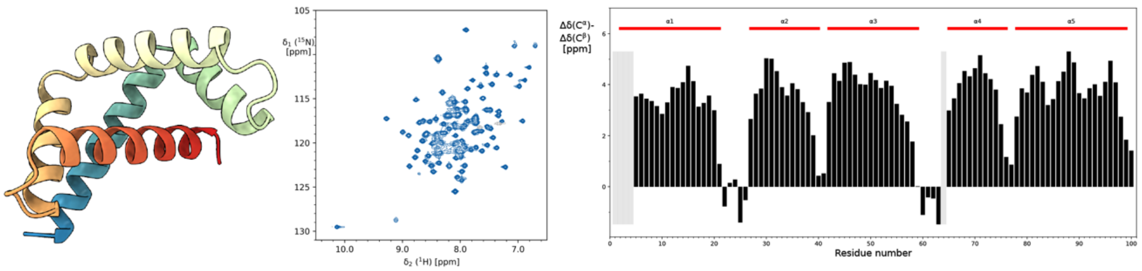


p3 B5

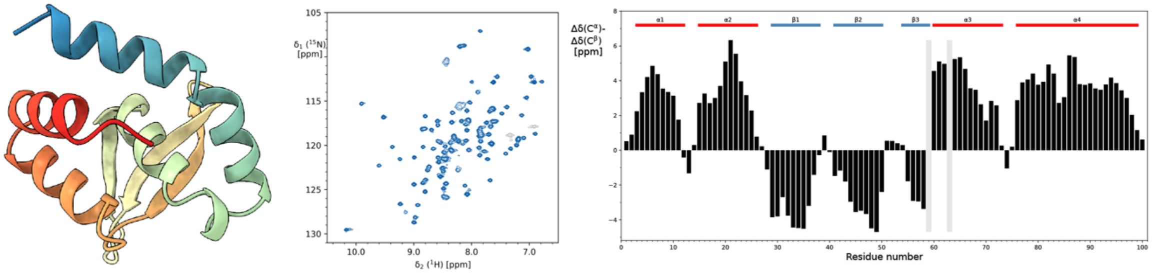


p3 E12

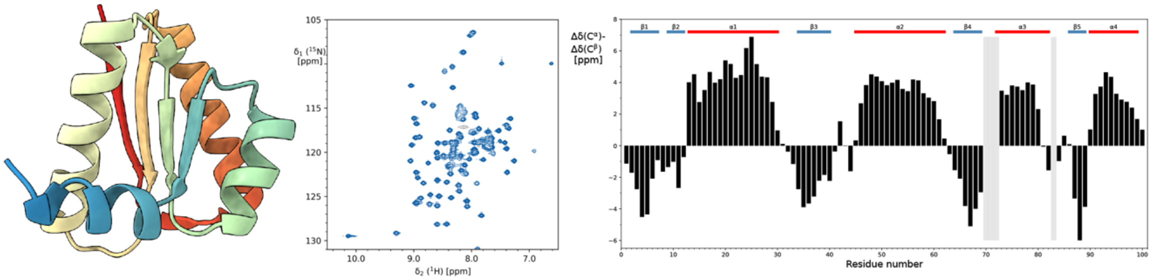

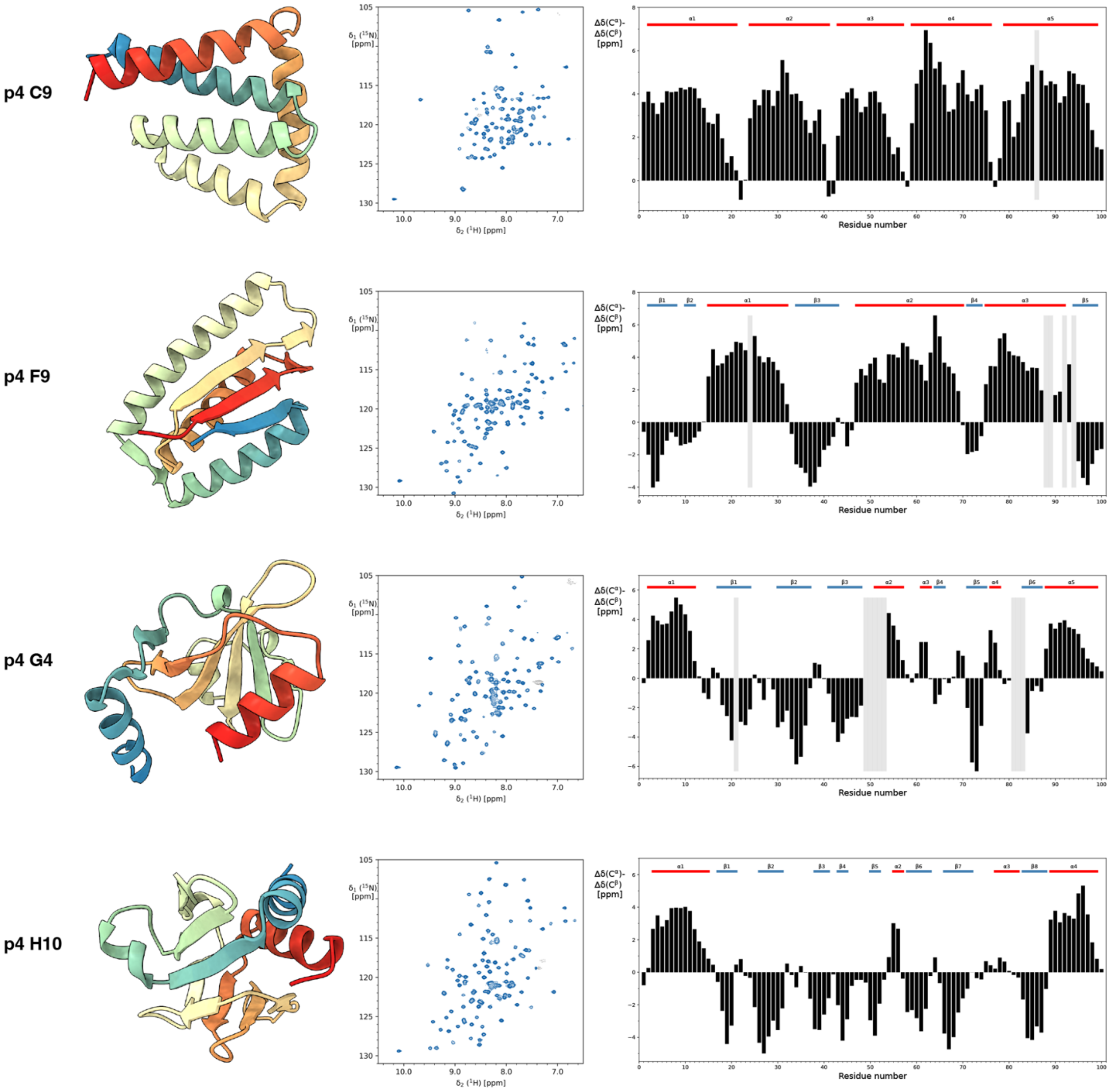


**Supplementary Figure S6.** Secondary structure plots of nine selected designer proteins. On the left, the designed structure; in the middle, the measured 2D [$^{15}$N,$^{1}$H]-SOFAST HMQC spectrum; and on the right, the secondary structure chemical shift plot. The expected secondary structure elements are indicated at the top, in red for α-helices and in blue for β-sheets.

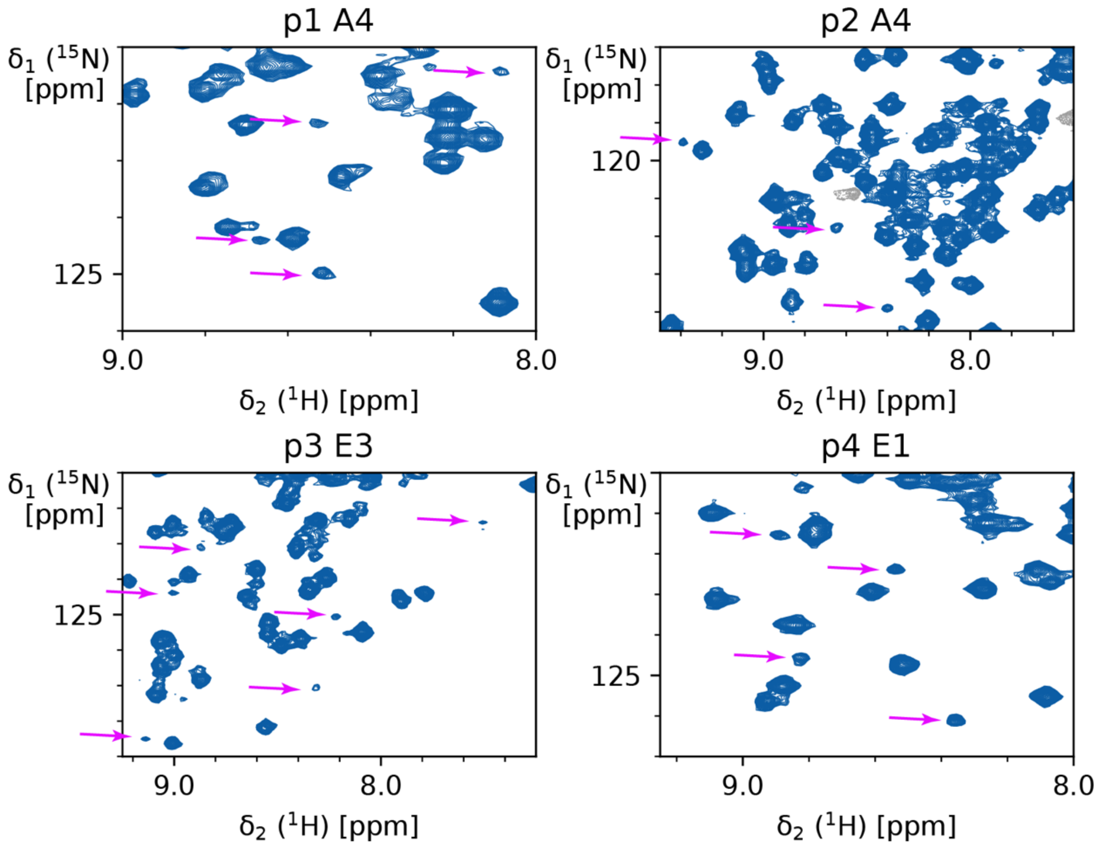


**Supplementary Figure S7.** Observation of high-energy states in designer proteins. Shown are enlarged regions from the 2D [$^{15}$N,$^{1}$H]-SOFAST HMQC spectra of four designer proteins. Minor peaks that likely correspond to high-energy conformations are indicated by purple arrows. These arise most likely from proline cis-trans isomerization.

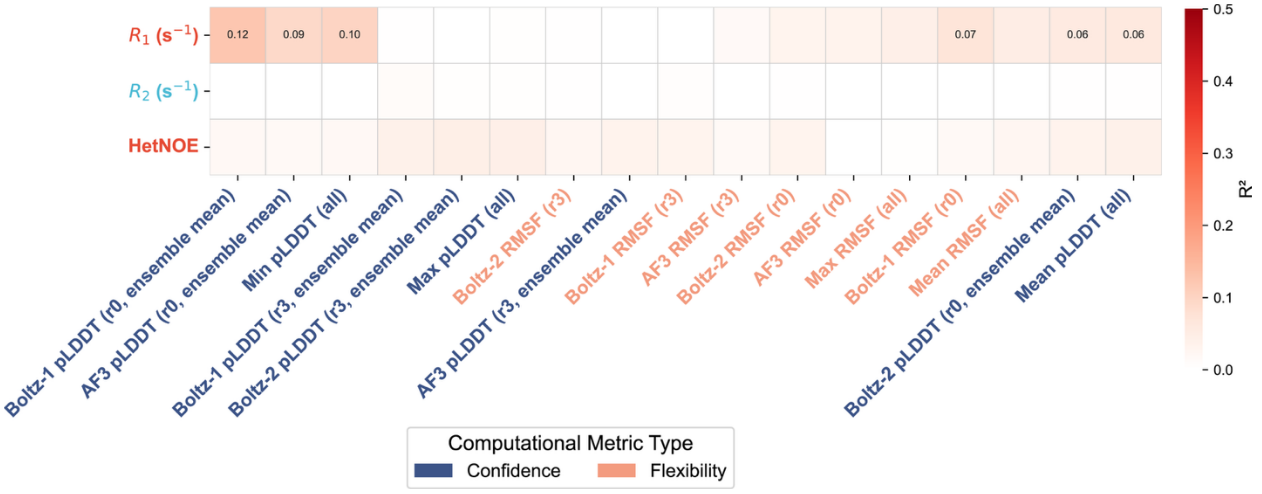


**Supplementary Figure S8.** Correlation between computational metrics and NMR relaxation parameters for 9 proteins. Computational metrics are shown on the x-axis, and color-coded by category. NMR relaxation parameters are shown on the y-axis. Coefficient of determination ($R^2$) are indicated as a colormap for each pair of computational-experimental metric. Coefficient greater than 0.1 are indicated explicitly as numerical values.
